# A comprehensive reference of mouse CD8αβ T cell differentiation states

**DOI:** 10.64898/2026.02.02.703365

**Authors:** Giovanni Galletti, Anna-Maria Globig, Olga Barreiro, Taylor A. Heim, Shuozhi Liu, Samantha M. Borys, Odhran Casey, Alexander Monell, Dhruv Patravali, Nicole E. Scharping, Sara Quon, Kennidy K. Takehara, Amir Ferry, Kitty P. Cheung, Ellen Duong, Tomoyo Shinkawa, Stefani Spranger, Samuel M. Behar, Susan M. Kaech, Ananda W. Goldrath, David Zemmour, the immgenT Project

## Abstract

Mouse CD8^+^ T cell differentiation has been studied extensively in models of infections and tumors, yet no unified framework spans the full spectrum of immunological contexts. Within the immgenT project, we profiled RNA, surface markers, and TCR clonotypes in conventional CD8^+^ T cells across >600 samples, spanning multiple perturbations, tissues, and timepoints. Twenty-one clusters across naive, effector, circulating memory, tissue-resident memory, progenitor-exhausted, and terminally-exhausted CD8^+^ T cell compartments emerged, with striking molecular convergence across acute and chronic infections, tumors, autoimmunity, aging, and homeostasis, illustrating that shared transcriptional states support protective or dysfunctional outcomes depending on developmental history and microenvironment. We validate immgenT as a comprehensive reference by integrating external datasets from conditions not represented in immgenT and by defining a flow cytometry panel spanning the CD8^+^ T cell landscape. Thus, immgenT-CD8 provides a molecular framework for harmonizing CD8^+^ T cell literature and clarifies relationships across diverse immune challenges.

## INTRODUCTION

CD8^+^ T cells are critical mediators of protective immunity against intracellular pathogens and tumors. Upon antigen encounter, naive CD8^+^ T cells undergo dramatic clonal expansion and acquire functional properties shaped by the intensity, duration, and anatomical location of stimulation^1^. In acute infections, most activated CD8^+^ T cells transiently adopt potent cytotoxic and migratory capabilities to clear the pathogen, while a fraction survives long-term with enhanced recall potential. Among long-lived CD8^+^ T cell populations, some recirculate through blood and lymphoid tissues to mount systemic responses upon re-challenge, whereas others permanently settle in non-lymphoid organs, providing immediate, localized protection at barrier and parenchymal sites such as skin, mucosa, liver, lung, and kidney^2^. These tissue-anchored cells exhibit rapid cytokine and cytotoxic responses upon local antigen re-encounter and are increasingly recognized as key correlates of durable vaccine– and immunotherapy-induced protection^3^. Their establishment depends on local environmental cues, including TGF-β, retinoic acid, and chemokine gradients^4,5^, which drive expression of integrins (e.g., CD103/αEβ7, CD49a/α1β1) and CXCR6^2^ for tissue retention and positioning. Transcriptional regulators including Hobit (*Zfp683*), Runx3, and others, promote and reinforce long-term residence, although cells in different organs rely on partially distinct gene modules^6–8^, reflecting microenvironmental adaptation and complicating the definition of universal residency signatures.

In settings of persistent antigen exposure, such as chronic viral infections or tumors, CD8^+^ T cells progressively lose effector functions, upregulate multiple inhibitory receptors (e.g., PD-1, TIM-3, LAG-3, TIGIT), and adopt altered metabolic states and transcriptional expression associated with TOX^9–13^. Within these populations, a subset retains stem-like self-renewal capacity, while the differentiated progeny have reduced cytokine-producing and cytotoxic potential^9,14–23^. The self-renewing compartment strongly influences the success of immune checkpoint blockade therapies^24–30^. Distinguishing long-term, tissue-anchored T cells arising after acute stimulation from those persisting under chronic antigen exposure remains challenging, as both share core residency programs and surface markers (e.g., CD103, CD49a, CXCR6), fueling debate about their developmental relationships and functional equivalence^31,32^.

Over the past two decades, the immunology community has categorized these context-dependent behaviors of CD8^+^ T cells into canonical subsets (e.g., circulating central-memory (T_CM_), effector-memory (T_EM_), tissue-resident memory (T_RM_), progenitor-exhausted (T_PEX_), and terminally-exhausted (T_EX_) CD8^+^ T cells), using a nomenclature that has proved invaluable for communication. However, widespread adoption of discrete labels has sometimes encouraged over-classification and obscured the molecular convergence evident from single-cell studies: shared transcriptional states can arise in dramatically different contexts and support protective or dysfunctional outcomes depending on developmental history and ongoing microenvironmental cues. This convergence, together with subtle laboratory variations in gating and naming of highly similar populations, has generated persistent ambiguity in the literature^33^. A unified, single-cell-resolution molecular reference capturing CD8^+^ T cell states across diverse perturbations, tissues, and time points could harmonize annotation and clarify biological relationships.

Here, we present the immgenT-CD8 framework, an integrated single-cell transcriptomic and surface-protein (128-marker panel) reference of >200,000 CD8αβ T cells from 678 samples spanning 80 experiments, 53 immunological challenges, and 45 tissues. By combining unbiased clustering, CITE-seq-derived combinatorial surface markers, and antigen-specific cell identification, we resolve 21 robust CD8^+^ T cell states that encompass naive, effector, circulating memory, tissue-resident memory, progenitor-exhausted, and terminally-exhausted populations. We define precise phenotypic signatures, flow cytometry gating strategies, interpretable gene-programs for each state, and uncover previously underappreciated heterogeneity. The web tool Rosetta2 allows users to interactively explore the atlas. With immgenT reference-based integration (T-RBI), we projected external CD8^+^ T cell datasets onto this unified map and validated that the immgenT-CD8 framework provides the first comprehensive single-cell coordinate system for mouse CD8^+^ T cell differentiation. This publicly accessible, curated resource (https://www.immgen.org/ImmGenT/) resolves fragmentation in the literature, offers immediately actionable tools for cell-state identification and isolation, and establishes a reference standard for mapping CD8^+^ T cell heterogeneity.

## RESULTS

### Scope and overview of CD8^+^ heterogeneity in immgenT

As part of the immgenT Open Source Project, we generated a comprehensive single-cell atlas of mouse conventional CD8αβ T cells. We profiled 206,160 CD8^+^ T cells across 678 samples from 80 independent experiments (IGT1-96), spanning 53 distinct immunological challenges (including acute and chronic infections, tumors, autoimmunity, aging, homeostasis) and 45 tissues (**Fig. 1a,b**, **Extended Data Table 1**). Of these cells, 21,292 were antigen-specific, identified by tetramer staining, congenic markers, or paired TCRαβ sequencing (e.g., P14 or OT-I cells; **Fig. 1b**). This dataset was generated using a standardized protocol that jointly measured gene expression (scRNA-seq) and 128 surface proteins (CITE-seq) with paired TCR sequencing (see **Methods**), as detailed in the companion immgenT-Cosmology manuscript^34^.

**Figure 1.**
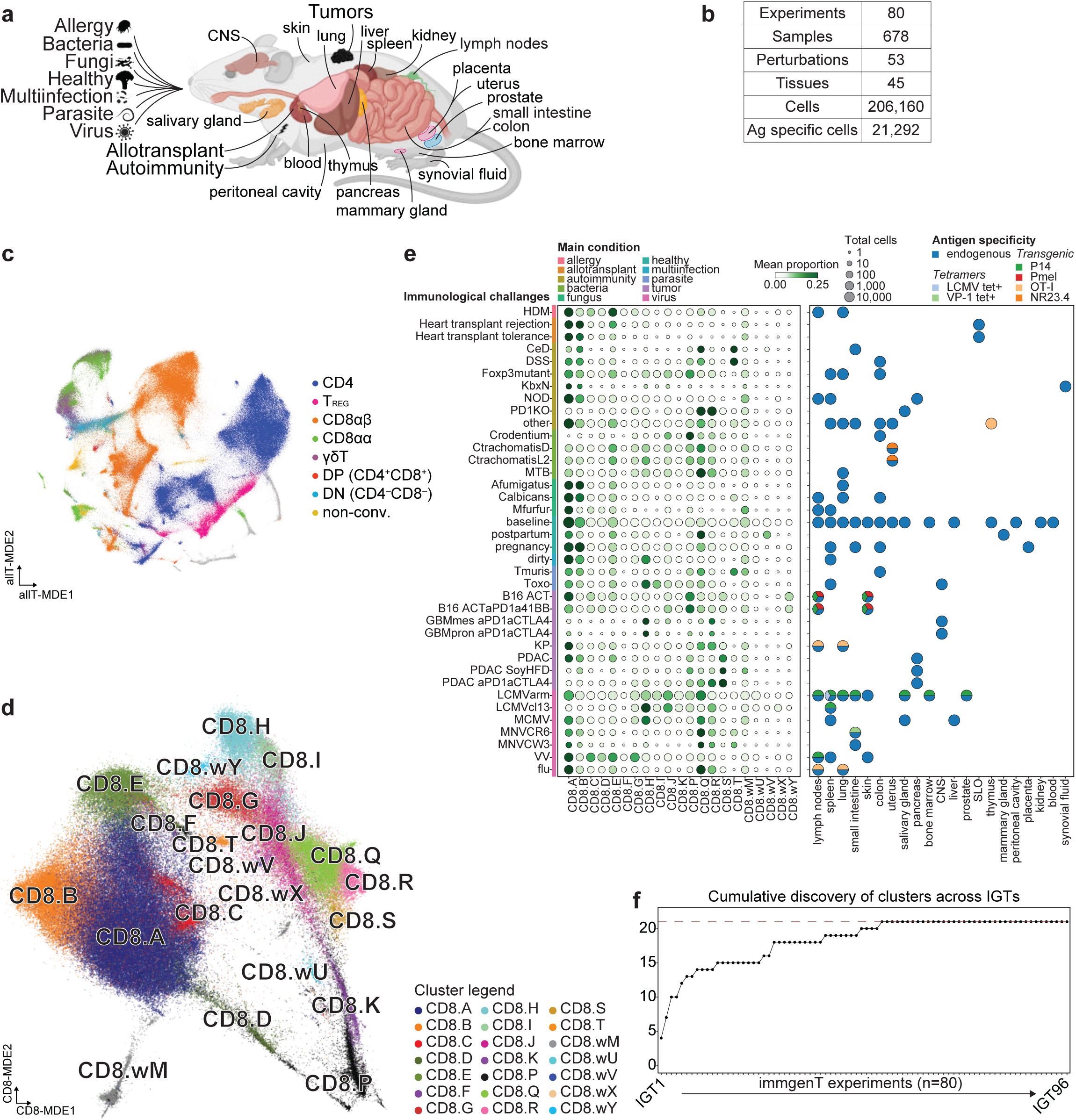
immgenT resolves CD8^+^ T cell heterogeneity into 21 discrete clusters and a reference embedding. **a**, Schematic of the conditions and main tissues selected to build the immgenT-CD8 framework. All experiments used single-cell RNAseq combined with 128-plex CITE-seq and TCR-seq; **b**, Table summarizing the main features of the total immgenT-CD8 dataset: 206,160 CD8αβ T cells profiled across 678 samples from 80 experiments (IGT1-96), spanning 53 immunological challenges (e.g., acute and chronic infections, tumors, autoimmunity, aging, homeostasis) and 45 tissues, including 21,292 antigen-specific cells identified by tetramer staining, congenic markers, or paired TCRαβ sequencing; **c**, Minimum Distortion Embedding of all T cells (allT-MDE) profiled in the immgenT dataset, colored by major T cell lineages: CD4^+^, CD8αβ, CD8αα, DN, DP, γδ T cells, T_REG_, and non-conventional, showing that CD8αβ T cells form a transcriptional continent separate from other lineages; **d**, CD8-MDE colored by cluster identity. Cluster labels prefixed with “w” denote provisional clusters, typically small, and are not further analyzed in this study; **e**, Overview of the dataset. For each immune condition (rows), the left dot plot shows the number of cells (dot size) and mean cluster proportion (color). The color scale goes from 0.00-0.25, where 0.25 indicates that the cluster accounts for 25% of all CD8^+^ T cells profiled in that condition category on average. The right dot plot indicates the profiled organs and whether antigen-specific cells were included in each immune condition: P14 transgenic TCR or LCMV tetramer^+^ identifies specificity for lymphocytic choriomeningitis virus; VP-1 tetramer^+^ for murine norovirus; NR23.4 for Chlamydia trachomatis; Pmel for B16 melanoma (gp100 tumor antigen); OT-I for ovalbumin antigen overexpressed in the models indicated (see **Extended Data Table 1**). Each condition includes samples with at least 10 total CD8^+^ T cells recovered in at least one cluster; **f**, Saturation of CD8^+^ clusters within the immgenT dataset. The number of clusters containing at least 100 cells across experiments reaches saturation after approximately half the experiments. **Abbreviations**: CNS, central nervous system; Ag, antigen; T_REG_, regulatory T cells; DP, double-positive; DN, double-negative; non-conv., non-conventional; tet+, tetramer^+^; HDM, house dust mite; CeD, celiac disease; DSS, dextran sulfate sodium; KbxN, K/BxN arthritis; NOD, non-obese diabetic; Crodentium, Citrobacter rodentium; CtrachomatisD, Chlamydia trachomatis serovar D; CtrachomatisL2, Chlamydia trachomatis serovar L2; MTB, Mycobacterium tuberculosis; Afumigatus, Aspergillus fumigatus; Calbicans, Candida albicans; Mfurfur, Malassezia furfur; Tmuris, Tabula muris; Toxo, Toxoplasma gondii; ACT, adoptive cell therapy; GBMmes, glioblastoma mesenchymal subtype; GBMpron, glioblastoma proneural subtype; KP, Kras^G12D^/p53^−/−^ lung adenocarcinoma; PDAC, pancreatic ductal adenocarcinoma; SoyHFD, soy high-fat diet; LCMVarm, LCMV-Armstrong; LCMVcl13, LCMV-Clone 13; MCMV, murine cytomegalovirus; MNVCR6, murine norovirus CR6; MNVCW3, murine norovirus CW3; VV, vaccinia virus; flu, influenza virus; SLOs, secondary lymphoid organs.

We used totalVI, a deep generative model that jointly models RNA and surface-protein counts to integrate all T cells in a shared latent space that was then projected in two dimensions using Minimum Distortion Embedding (MDE) (**Fig. 1c**). Within the allT-MDE, CD8αβ T cells clustered separately from the other 7 lineages (**Fig. 1c** and **Extended Data Fig. 1a**). To resolve further CD8^+^ population structure, we also generated a CD8^+^ specific dimensionality reduction plot (CD8-MDE) and identified 21 shared clusters (**Fig. 1d**), recapitulating the CD8^+^ T cell heterogeneity derived from the immunological challenges and tissues analyzed (**Fig. 1e** and **Extended Data Table 2**). The number of clusters saturated after approximately half of the experiments (**Fig. 1f**). The immgenT cluster annotation is harmonized across the T cell lineages (CD8, CD4, T_REG_, etc.), so that similar states use the same alphabet letter^34^ (e.g., P always denotes clusters of proliferating cells); to achieve this some letters are not used. Provisional clusters of unclear significance were annotated with a ‘w’ prefix (namely ‘work in progress’). Among them, the CD8.wM state corresponds to a shared transcriptional state observed across multiple T cell lineages; its biological significance remains unclear and is discussed in the immgenT-Cosmology manuscript^34^. We validated the immgenT reference by projecting eight independent published CD8^+^ T cell datasets (104,024 cells total) onto the map using T-RBI, our reference-based integration method (**Extended Data Fig. 1b,c** and **Extended Data Table 3**, see **Methods**). Projection accuracy was high for all datasets, and no novel states emerged outside the existing 21 immgenT clusters.

To orient the atlas within classical immunological terminology, we examined CD62L and CD44 protein levels (CITE-seq). This stratification divided cells into the expected broad compartments including naive/resting (CD62L^+^CD44^−^), effector/effector-memory-like (CD62L^−^CD44^+^), central-memory-like (CD62L^+^CD44^+^), and double-negative (CD62L^−^CD44^−^) cells; each compartment contained multiple transcriptionally distinct clusters (**Extended Data Fig. 1d**). The 21 clusters presented dynamic gene– and protein-expression profiles (**Fig. 2a**). Classic subset-defining genes (e.g., *Tox*, *Pdcd1*, *Cxcr6*, *Tcf7*) were expressed more broadly than anticipated, underscoring that single transcripts or small panels are insufficient to define states. The 21 CD8^+^ T cell states were detectable widely across conditions, but their relative abundances shifted dramatically depending on the challenge, even at healthy baseline (**Fig. 2b,c** and **Extended Data Fig. 1e**). In line with their likely antigen-encounter history, CD8^+^ T cells from clusters CD8.G-S showed a high degree of clonotype sharing and a more pronounced clonal expansion than the resting states CD8.A-F (**Extended Data Fig. 1f,g**).

**Figure 2.**
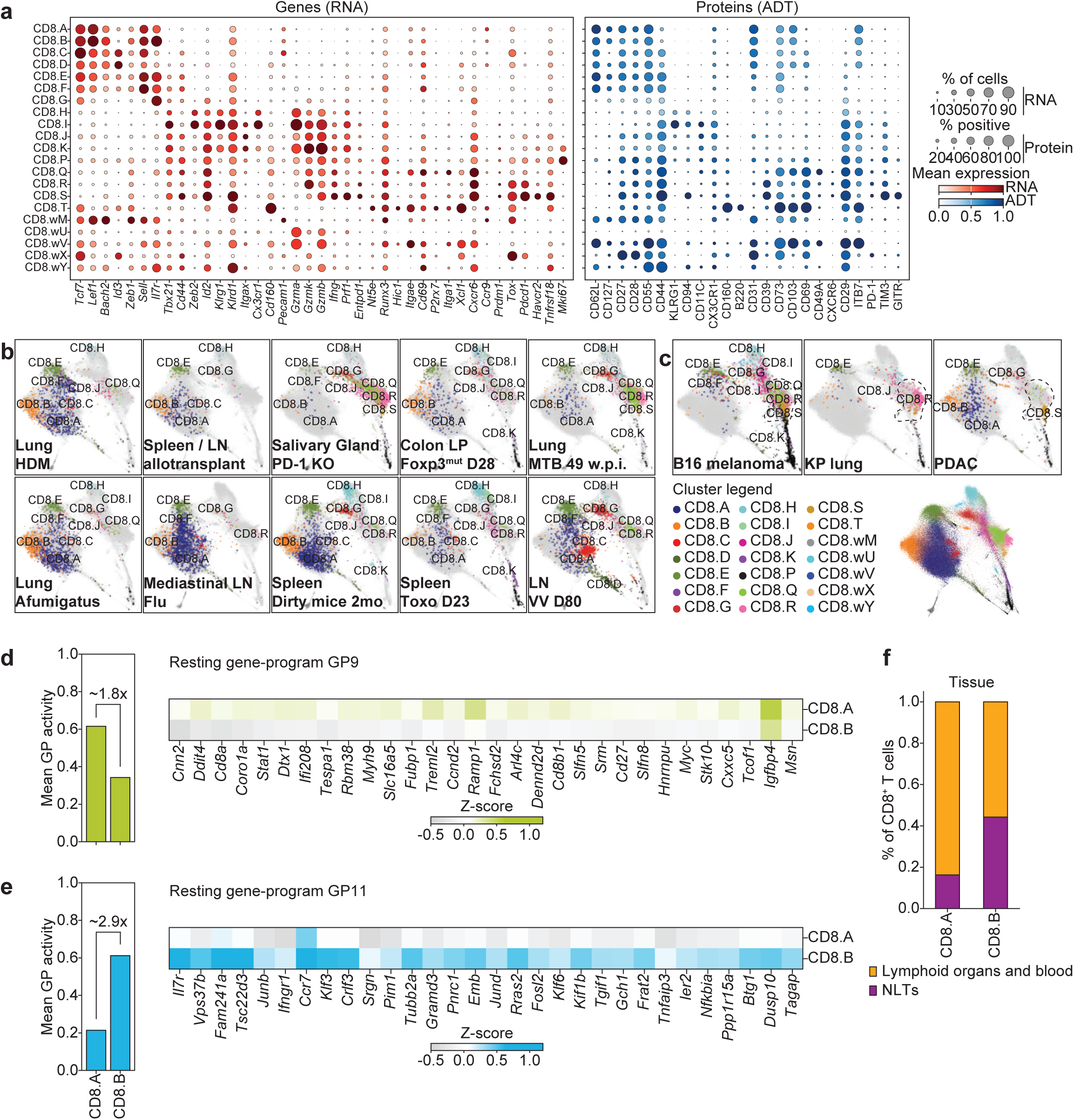
Shared CD8^+^ T cell states across immune conditions. **a**, Dot plots showing expression of common CD8^+^ T cell differentiation genes (left) and surface markers (right) across clusters. Color indicates scaled mean expression, and dot size represents the fraction of cells expressing each gene or protein; **b,c**, CD8-MDE highlighting selected experiments illustrating that the same 21 molecular states recur across different immunological challenges with condition-specific shifts in abundance. Cells are colored by cluster identity; background immgenT-CD8 are shown in gray; **d,e**, Bar plots showing the mean gene-program (GP) activity of resting GP9 (**d**) and GP11 (**e**) in cells from clusters CD8.A and CD8.B (left) and heatmaps of the top 30 genes (Z-score normalized) from each GP (right). GPs were derived by empirical Bayes matrix factorization of the full dataset; **f**, Stacked bar plot showing the fraction of cells from clusters CD8.A and CD8.B that belong to lymphoid organs and blood versus non-lymphoid tissues (NLTs); lymphoid organs include spleen, thymus, lymph nodes, bone marrow; NLT comprised all remaining non-tumor, non-lymphoid tissues. Tumor samples were excluded from this analysis. **Abbreviations**: ADT, Antibody-Derived Tag; HDM, house dust mite; LN, lymph node; KO, knock-out; LP, lamina propria; w.p.i., weeks post-infection; mo, months; D, day; VV, vaccinia virus; GP, gene-program; n., number.

We next highlighted well-studied models directly on the reference map. In acute LCMV-Armstrong and chronic LCMV-Clone 13 infections, days 7-8 splenic effector CD8^+^ T cells converged in clusters CD8.I, CD8.J, and CD8.K, while memory time points (days 27-30+) diverged based on infection: LCMV-Armstrong enriched clusters CD8.E, CD8.F, CD8.G, and CD8.H, whereas LCMV-Clone 13 enriched clusters CD8.H and CD8.R (**Extended Data Fig. 1h**). In non-lymphoid tissues such as the small intestine, effector CD8^+^ T cells transitioned predominantly into cluster CD8.Q (**Extended Data Fig. 1h**). These patterns held for both endogenous and antigen-specific cells (although at late timepoints, after resolution of inflammation, the baseline population of resting splenic T cells was apparent in the endogenous compartment). Thus, the identified cell states robustly capture antigen-driven differentiation programs and are not merely dominated by endogenous bystander features.

In other immunological contexts, we detected fewer clusters with stronger dominance of one or two populations, whereas in others we observed mixtures of clusters that, in LCMV infections, were typically associated with distinct time points or tissues. Clusters CD8.I, CD8.J, and CD8.K, associated with acute LCMV responses, were also present in autoimmune models such as scurfy/Foxp3-deficient mice, yet here they appeared together with clusters that were not normally co-enriched during LCMV infection (**Fig. 2b**). Clusters CD8.Q, CD8.R, and CD8.S were enriched in chronic LCMV-Clone 13 and cancer settings (**Fig. 2c**). Even within tumors, such as B16 melanoma, broader heterogeneity emerged, including contributions from clusters CD8.E, CD8.F, CD8.I, CD8.J, CD8.K (**Fig. 2c**). Finally, nearly all clusters were detectable, albeit some at low frequencies, in the tissues of unchallenged baseline mice (**Extended Data Fig. 1e**). Together, these observations suggest that CD8^+^ T cell clusters may be more pleiotropic and more flexibly deployed across a range of immune contexts.

The naive CD8^+^ T cell compartment showed subtle heterogeneity. CD62L^+^CD44^−^ cells mapped overwhelmingly to clusters CD8.A and CD8.B and expressed canonical naive T cell-associated genes (*Sell*, *Ccr7*, *Tcf7*) (**Fig. 2a** and **Extended Data Fig. 1d**). For further insights, we applied gene-program (GP) analysis, an empirical Bayes matrix factorization that deconstructs transcriptomes into compact, biologically interpretable programs (see **Methods** and companion immgenT-Cosmology manuscript^34^). CD8.A showed stronger activity of gene-program GP9 and originated predominantly (>80%) from lymphoid organs and blood, whereas CD8.B showed stronger gene-program GP11 activity and contained a higher fraction (∼50%) of cells from non-lymphoid tissues (**Fig. 2d-f**). Notably, gene-expression signatures of previously defined CD5^HI^ versus CD5^LO^ naive CD8^+^ T cells were modestly enriched in clusters CD8.A and CD8.B, respectively (**Extended Data Fig. 1i**), in line with published studies of naive T cell heterogeneity^35,36^.

Together, these analyses show that the immgenT-CD8 reference captures a broad spectrum of differentiation from naive to effector, memory, resident, and exhaustion-like CD8^+^ T cell states. The 21 established clusters represent recurrent transcriptional and proteomic programs that are deployed across time, tissues, and diverse immune challenges rather than being restricted to any single context. We next examine these states in detail, beginning with the effector, circulating memory, tissue-resident memory, and exhaustion-like CD8^+^ T cell populations in canonical LCMV and tumor models before extending to broader immunological settings.

### Transcriptional, clonal, and phenotypic heterogeneity of effector CD8^+^ T cells

At the peak of expansion (days 7-8) after both acute LCMV-Armstrong and chronic LCMV-Clone 13 infection, antigen-specific CD8^+^ T cells mapped predominantly to clusters CD8.I, CD8.J, and CD8.K (**Fig. 3a** and **Extended Data Fig. 2a**). Typical analyses of effector T cell populations (i.e., day 7 of infection) subdivide by cell-surface expression of KLRG1 and CD127. At the peak of the CD8^+^ T cell response, antigen-specific CD8^+^ T cells include an “early-effector” KLRG1^−^CD127^−^ population^37,38^, which can give rise to both the KLRG1^+^CD127^−^ terminally-differentiated, short-lived effector and KLRG1^−^CD127^+^ memory-precursor cell populations^1,39,40^. In addition, a KLRG1^+^CD127^+^ population with intermediate T-bet expression, reduced IL-2 production, and retained proliferative potential has been described^41^. Standard KLRG1/CD127 gating can leave many CD127^−^ KLRG1-low/negative cells unaccounted for, underestimating the full diversity of the short-lived effector cell compartment.

**Figure 3.**
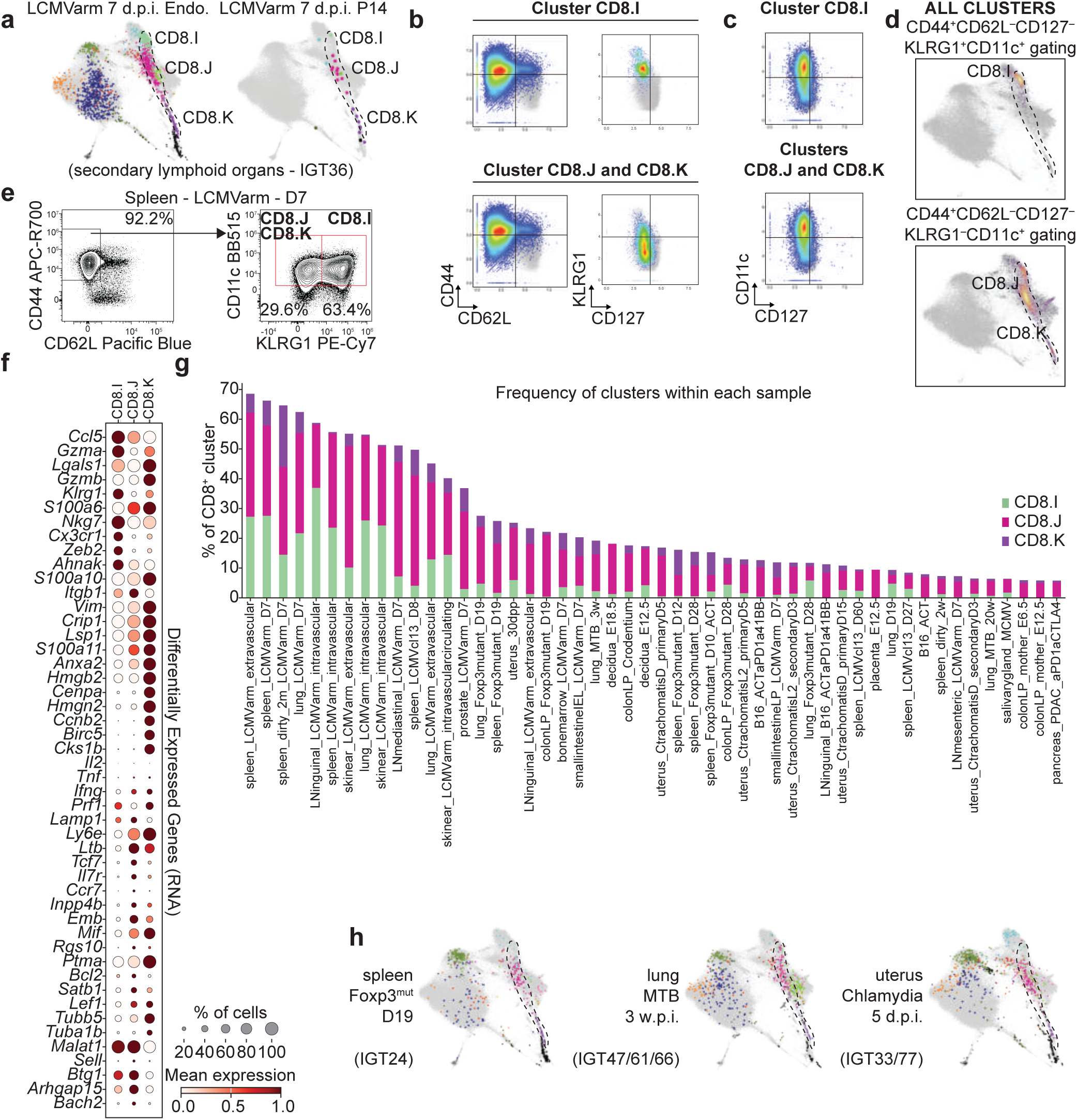
Effector CD8^+^ T cells comprise three distinct recurrent states. **a**, CD8-MDE showing endogenous (left) and LCMV-specific (P14) T cells (right) at the peak effector phase (days 7-8 post-infection). Cells are colored by cluster identity; background immgenT-CD8 are shown in gray; **b**, Scatter plots showing predicted CITE-seq-based (log1p(CP10K)) gating strategies for isolation of the indicated clusters using surface proteins (CD44, CD62L, CD127, KLRG1); **c**, Scatter plots showing CITE-seq (log1p(CP10K)) protein expression of CD11c and CD127 on cells from clusters CD8.I, CD8.J, and CD8.K (density) versus all other CD8^+^ T cells (gray). CD11c shows 94% specificity for CD8.I, .J, and .K versus all other clusters (**Extended Data Table 4**); **d**, Gating enrichment projection onto the CD8-MDE using the CD44^+^CD62L^−^CD127^−^CD11c^+^ strategy (with KLRG1 for further subdivision), showing tight localization to the CD8.I/CD8.J/CD8.K region; **e**, Representative flow cytometry plots showing CD44, CD62L, CD11c, and KLRG1 expression in splenic CD8^+^ T cells at day 7 after LCMV-Armstrong infection (percentages shown in gates); **f**, Dot plot showing expression of a curated list of differentially expressed genes across clusters CD8.I, CD8.J, and CD8.K. Color indicates scaled mean expression, and size represents the fraction of cells expressing each gene; **g**, Stacked bar plot showing the proportion of clusters CD8.I, CD8.J, and CD8.K within the top 50 samples. Only samples with at least 10 total CD8^+^ T cells recovered in at least one cluster are shown (sample description details in **Extended Data Table 1**); **h**, CD8-MDE showing CD8^+^ T cells in the spleen of 19-day-old Foxp3-deficient mice (left), in the lung of MTB-infected mice 3 weeks post-infection (middle) and in the uterus 5 days post-infection with C. trachomatis. **Abbreviations**: d.p.i., days post-infection; Endo., endogenous; D, day; w.p.i., weeks post-infection; MTB, Mycobacterium tuberculosis.

In the immgenT framework, CITE-seq surface-protein data showed that clusters CD8.I, CD8.J, and CD8.K were CD44^+^CD62L^−^CD127^−^, with only cluster CD8.I expressing detectable KLRG1 protein (**Fig. 3b**). In contrast, CD11c protein was expressed at high levels across all three clusters (specificity 94% relative to all other CD8^+^ T cell states; **Extended Data Table 4**) (**Fig. 3c**). Thus, we propose that the combined phenotype CD44^+^CD62L^−^CD127^−^ CD11c^+^ captures the full short-lived effector CD8^+^ T cell populations, while KLRG1 further distinguishes cluster CD8.I from CD8.J and CD8.K. To further illustrate this gating strategy, we applied it to the reference map and confirmed strong enrichment in the CD8.I/CD8.J/CD8.K region of the MDE (**Fig. 3d**). Independent flow cytometry validation on day 7 LCMV-Armstrong splenic CD8^+^ T cells confirmed CD11c expression on CD62L^−^CD44^+^ cells, with only a subset co-expressing KLRG1 (**Fig. 3e**). CD11c levels were high specifically during the acute effector phase, while low on naive and memory T cell populations (**Extended Data Fig. 2b,c**). Using this approach required bright fluorophores with careful titration because expression on T cells is lower than on dendritic cells (**Extended Data Fig. 2d**). Time-course flow cytometry analysis aligned with published scRNA-seq data showing *Itgax* transcript peaking at day 6 (**Extended Data Fig. 2e,f**).

These three clusters therefore represent, at transcriptional and surface-protein (CITE-seq panel) level, distinct states within the canonical short-lived effector compartment. While functional differences among them remain to be fully tested, the data show that the CD127^−^ effector pool classically viewed as relatively homogeneous or only subdivided by KLRG1 contains at least three recurrent molecular states. CD11c provides a more inclusive surface marker than KLRG1 for isolating this compartment across infections and tissues. To relate immgenT clusters to classical memory-precursor effector cells (MPECs), we examined the canonical MPEC phenotype (KLRG1^−^CD127^+^) within the CD44^+^CD11c^+^ gate. A small fraction of cells with this phenotype mapped to cluster CD8.G (discussed in the next section), while the majority of the CD127^−^ effector CD8^+^ T cell compartment fell into clusters CD8.I, CD8.J, and CD8.K (**Extended Data Fig. 2g**). Thus, in the immgenT framework, classical MPECs do not form a fully separate cluster at the peak of expansion but are embedded within the broader effector-to-memory continuum, with cluster CD8.G showing partial overlap.

Within the effector CD8^+^ T cell populations, gene expression was heterogeneous (**Fig. 3f**). Many differentially expressed transcripts belonged to gene-programs GP10 and GP25 (**Extended Data Fig. 2h,i**), which are also active in cluster CD8.H (a terminally-differentiated effector-memory/long-lived effector state described in the next section). TCR clonal analysis revealed substantial sharing among clusters CD8.I, CD8.J, and CD8.K, consistent with branched differentiation or interconversion from common progenitors (**Extended Data Fig. 2j**). Across the full atlas, these clusters were not only enriched in acute LCMV-Armstrong responses (both lymphoid and non-lymphoid tissues) but also appeared at lower frequencies in other infections (e.g., Mycobacterium tuberculosis in the lung, Chlamydia in the uterus), autoimmunity (Foxp3-deficient mice), and tumors, particularly under checkpoint blockade or adoptive cell transfer (**Fig. 3g,h**), illustrating the broad use of these molecular states during CD8^+^ T cell immune responses.

In summary, clusters CD8.I, CD8.J, and CD8.K represent a family of interconnected but molecularly distinct states within the short-lived effector CD8^+^ T cell compartment. These states recur across multiple infections, autoimmunity, and tumor settings, and the cell-surface phenotype CD44^+^CD62L^−^CD127^−^CD11c^+^ provides a practical tool for their identification and isolation by flow cytometry.

### Heterogeneous recirculating memory CD8^+^ T cell subsets are broadly present across immune challenges and tissues

Clusters CD8.E, CD8.G, and CD8.H (and to a lesser extent CD8.F) dominate the late/memory phase (days 27-30+) of antigen-specific CD8^+^ T cell responses in secondary lymphoid organs after acute LCMV-Armstrong infection (**Fig. 4a**). A fraction of tetramer^+^ cells at memory time points mapped to naive-like clusters CD8.A and CD8.B, likely reflecting low-avidity or non-specific binding. Projection of an independent public dataset^6^ (P14 cells from blood and spleen at day 32 after LCMV-Armstrong) using T-RBI also confirmed that these clusters correspond to classical circulating memory populations (**Fig. 4b**).

**Figure 4.**
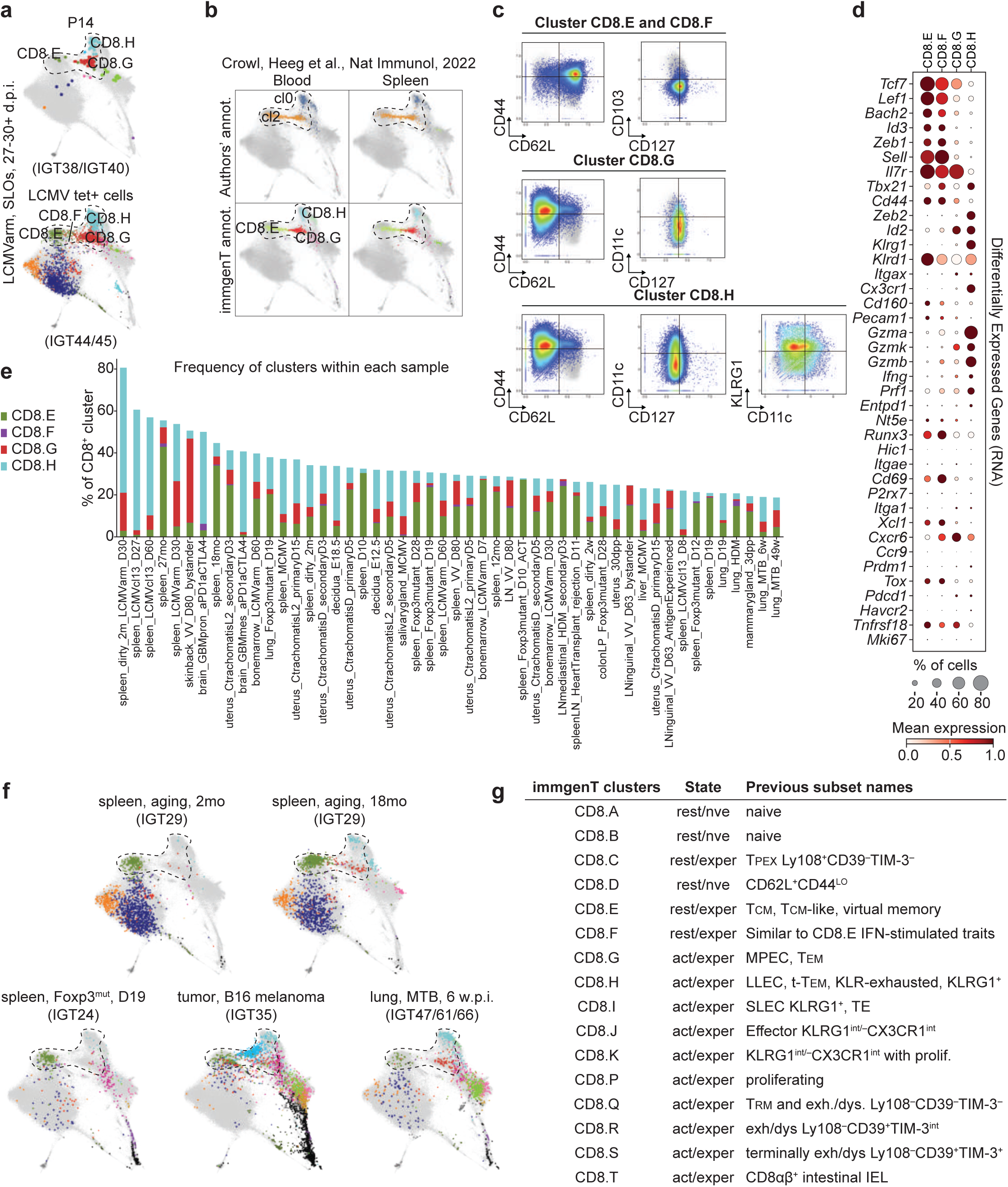
Circulating memory CD8^+^ T cells map to three main clusters that are present broadly beyond acute infections. **a**, CD8-MDE showing LCMV-specific T cells (P14 transgenic cells, top; GP33-tetramer^+^ cells, bottom) at late memory time points (days 27-30+) after acute LCMV-Armstrong infection in secondary lymphoid organs. Cells are colored by cluster identity; background immgenT-CD8 are shown in gray; **b**, T-RBI (immgenT reference-based integration) of published datasets of P14 CD8^+^ T cells from blood (left) and spleen (right) at day 32 after LCMV-Armstrong from Crowl et al.^6^). Cells are colored by authors’ annotation (top) and immgenT cluster identity (bottom); background immgenT-CD8 are shown in gray; **c**, Scatter plots showing CD62L, CD44, CD127, KLRG1 expression in clusters CD8.E, .F, .G and .H (CITE-seq; log1p(CP10K)); **d**, Dot plot showing scaled mean expression of common memory– and effector-associated genes across clusters CD8.E-H; **e**, Stacked bar plot showing the proportion of clusters CD8.E, CD8.F, CD8.G, and CD8.H within the top 50 samples. Only samples with at least 10 total CD8^+^ T cells recovered in at least one cluster are shown (sample description details in **Extended Data Table 1**); **f**, CD8-MDE showing examples of conditions in which CD8.E-H are present beyond acute LCMV-Armstrong: in the spleens of 2-month-old and 18-month-old mice, in the spleen of 19-day-old Foxp3-deficient mice, in B16 melanoma tumors, and in the lung of MTB infected mice (6 weeks post infection); **g**, Table showing the correspondence between the immgenT clusters, T cell state, and previous subset names. Provisional clusters (CD8.wM-Y) have been excluded from this table (see **Extended Data Table 5**). **Abbreviations**: d.p.i., days post-infection; tet, tetramer; annot., annotation; cl, cluster; mo, months; MTB: Mycobacterium tuberculosis; w.p.i., weeks post-infection; D, day; rest, resting; nve, naive; exper, antigen-experienced; act, activated; int, intermediate; exh., exhausted; dys., dysfunctional; IEL, intraepithelial lymphocytes.

Surface-protein profiles (CITE-seq and flow cytometry) aligned with classical delineations^42–45^ (**Fig. 4c** and **Extended Data Fig. 3a**): clusters CD8.E and CD8.F were CD62L^+^CD44^+^CD127^+^ (T_CM_); cluster CD8.G was CD62L^−^ CD44^+^CD127^+^ (T_EM_); and cluster CD8.H was CD62L^−^CD44^+^CD127^−^ with elevated KLRG1 (often termed long-lived effector cells, LLEC, or terminally-differentiated effector-memory, t-T_EM_). Previous studies^43,45^ defined LLEC as a persistent memory subset with effector-like features (high KLRG1, granzyme B, CX3CR1), homeostatic proliferation, and tissue-entry flexibility upon rechallenge. Similarly, a distinct t-T_EM_ subset was also described^44^ as a KLRG1^+^ terminally-differentiated population with potent cytotoxicity and limited multipotency/recall. Despite contextual differences, these populations share core phenotypes (high KLRG1, low CD127, low CD62L) and transcriptional signatures, suggesting overlapping definitions. These phenotypes share core features with the “exhausted KLR” state described in chronic LCMV and tumors^46,47^, highlighting molecular convergence across settings of acute and chronic antigen exposure. CITE-seq protein-based gating mapped these phenotypes to their clusters (**Extended Data Fig. 3b**), with independent flow cytometry confirming matching populations in day 30 LCMV-Armstrong endogenous CD8^+^ T cells (**Extended Data Fig. 3a**). Importantly, the transcriptional and surface-protein features analyzed here do not by themselves distinguish protective from dysfunctional states and functional studies will be required to resolve context-specific roles.

Gene expression followed expected patterns: clusters CD8.E, CD8.F, and CD8.G expressed variable levels of memory-associated genes such as *Tcf7*, *Id3*, and *Il7r*. In contrast, CD8.H expressed *Zeb2*, *Cx3cr1*, lacked *Il7r*, and maintained high levels of effector genes (*Gzma*, *Gzmb*, *Gzmk*, *Ccl5*, *Klrg1*) (**Fig. 4d**). Cluster CD8.H was enriched for gene-program GP10, which is also active in the acute effector clusters CD8.I/J/K (**Extended Data Fig. 2h**). Cluster CD8.F showed enrichment for GP16, an interferon-responsive program^48,49^ (e.g., *Ifit1*, *Ifit3*, *Isg15*), distinguishing it from cluster CD8.E (**Extended Data Fig. 3c**).

Importantly, these states are not restricted to LCMV-Armstrong (**Fig. 4e,f**). Clusters CD8.E-H appeared in secondary lymphoid organs during other infections, in non-lymphoid tissues (e.g., uterus after Chlamydia, lung after Mycobacterium tuberculosis), in tumor-infiltrating lymphocytes, in autoimmune models (Foxp3-deficient mice), and even at baseline homeostasis. Clusters CD8.E and CD8.F increased with aging in SPF mice, consistent with virtual memory-like cells^50^. Bystander-activated CD8^+^ T cells (e.g., after vaccinia scarification^51^) also mapped to clusters CD8.E-H (**Extended Data Fig. 3d**). Cluster-defining gene signatures remained stable across these diverse conditions (**Extended Data Fig. 3e-h**). Thus, the circulating memory CD8^+^ T cell subsets classically described as T_CM_, T_EM_, and t-T_EM_/LLEC map to clusters CD8.E, CD8.F, CD8.G, and CD8.H in the immgenT framework (**Fig. 4g and Extended Data Table 5**). However these molecular states recur across acute infections, tumors, autoimmunity, aging, and homeostasis, including settings of persistent antigen where “true” memory cannot form. These results caution against labeling all cells in these clusters, or with these shared phenotypic patterns, strictly as T_CM_, T_EM_, or t-T_EM_/LLEC, as the same transcriptional programs may be deployed in multiple immunological contexts.

### A high-resolution molecular framework of tissue-resident memory CD8^+^ T cells in non-lymphoid organs

Cluster CD8.Q emerged as the dominant state for antigen-specific CD8^+^ T cells in non-lymphoid tissues at memory time points. For example, CD8.Q contained the majority of P14 cells from non-lymphoid tissues at day 30+ post LCMV-Armstrong infection (**Fig. 5a-c**) and OT-I cells in the lung on day 23 of flu-OVA infection (**Fig. 5d**). Projection of an independent public dataset^6^ (P14 cells from small intestine and kidney on day 32 of LCMV-Armstrong) using T-RBI confirmed that canonical tissue-resident memory (T_RM_) cells across barrier and parenchymal tissues map predominantly to this single cluster (**Fig. 5e**).

**Figure 5.**
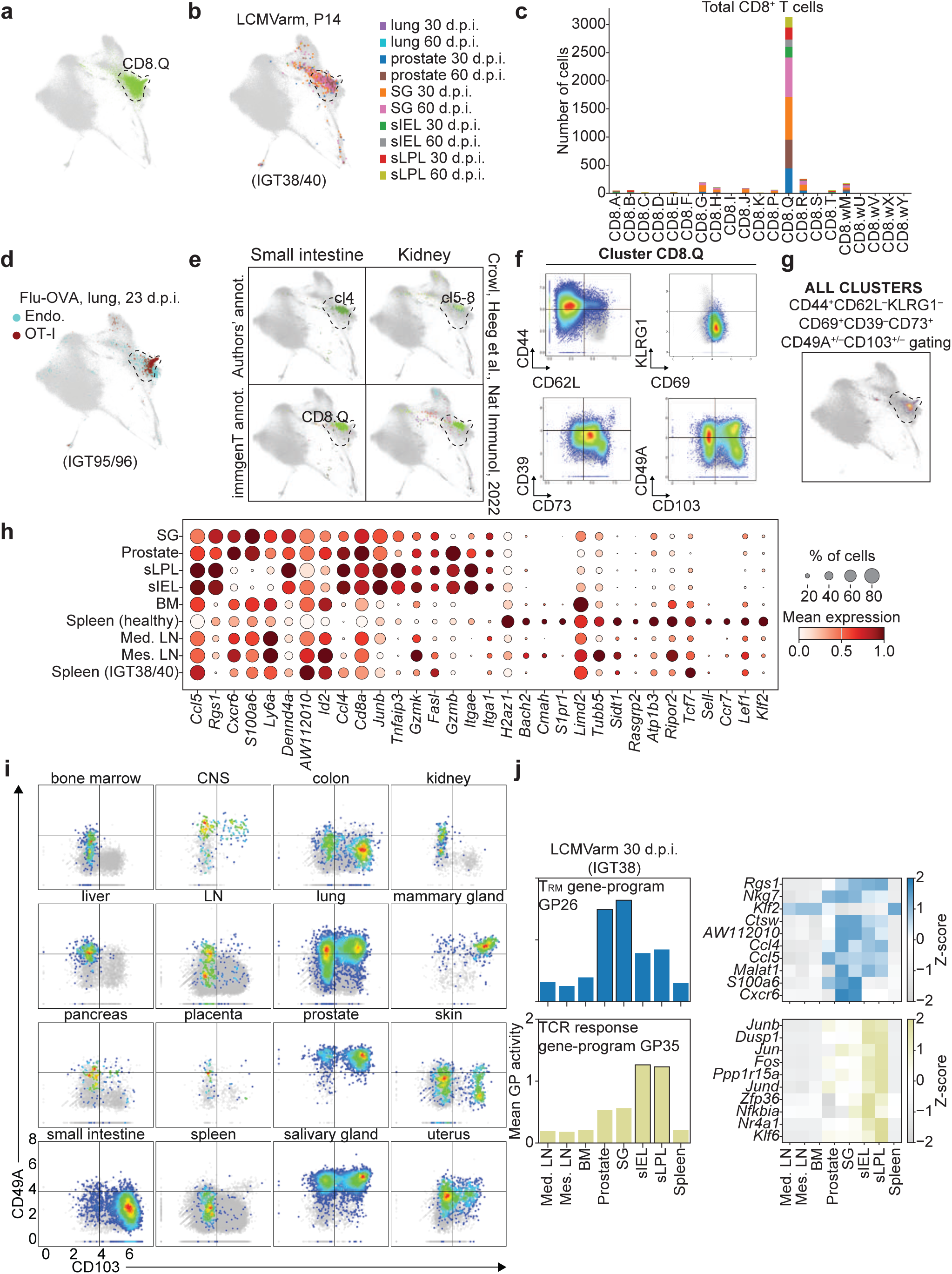
Tissue-resident memory CD8^+^ T cells converge on a single dominant transcriptional state while maintaining tissue-specific differences. **a**, CD8-MDE highlighting CD8.Q in green; **b,c**, CD8-MDE (**b**) and stacked bar plot (**c**) showing LCMV-specific T cells (P14 transgenic cells) at late memory time points (days 27-30+) after LCMV-Armstrong infection in non-lymphoid tissues (NLTs). Cells are colored by organ; background immgenT-CD8 are shown in gray; **d**, CD8-MDE showing flu-specific (OT-I) and endogenous T cells at day 23 post flu-OVA infection in the lung; **e**, T-RBI (immgenT reference-based integration) of published datasets of P14 CD8^+^ T cells from small intestine (left) and kidney (right) at day 32 after LCMV-Armstrong from Crowl et al.^6^. Cells are colored by authors’ annotation (top) and immgenT cluster identity (bottom); background immgenT-CD8 are shown in gray; **f**, Scatter plots showing predicted CITE-seq (log1p(CP10K)) surface-protein gating strategies for cluster CD8.Q (CD44, CD62L, CD69, CD73, CD39, KLRG1, and CD103/CD49a); **g**, CD8-MDE highlighting cells bioinformatically gated using the most optimal gating strategy for CD8.Q: CD62L^−^CD44^+^CD69^+^KLRG1^−^, CD39^−^CD73^+^, CD103^+^ or CD49A^+^; **h**, Dot plot showing scaled mean expression of CD8.Q-signature genes across tissues from CD8^+^ T cells at day 30+ after LCMV-Armstrong; **i**, Scatter plots showing CD103 vs CD49A expression in clusters CD8.Q across tissues (CITE-seq; log1p(CP10K)); **j**, Bar plots showing the mean gene-program (GP) activity of T_RM_ GP26 (top) and TCR-response GP35 (bottom) in CD8.Q cells at day 30+ after LCMV-Armstrong (left) and heatmaps of the top 10 genes (Z-score normalized) from each GP (right). **Abbreviations**: d.p.i., days post-infection; Endo., endogenous; annot., annotation; cl, cluster; CNS, central nervous system; LN, lymph node; GP, gene-program; Med., mediastinal; Mes., mesenteric; SG, salivary gland; sIEL, small intestine epithelium; sLPL, small intestine lamina propria.

CITE-seq surface-protein analysis within cluster CD8.Q showed consistent expression of CD44, CD69, and CD73, with absence of CD62L, KLRG1, and CD39 (**Fig. 5f**). These CITE-seq-guided gates mapped tightly back to CD8.Q on the MDE (**Fig. 5g**). Cluster CD8.Q signature genes were conserved across tissues (even lymphoid) at late LCMV-Armstrong time points and overlapped extensively with previously published T_RM_ signatures^6,52,53^ (**Fig. 5h** and **Extended Data Fig. 4a**). However, expression of the classic integrins associated with tissue residency, CD103 (αEβ7) and CD49a (α1β1), was heterogeneous within cluster CD8.Q (**Fig. 5f-i**). CD103 expression predominated for CD8^+^ T cells isolated from the small intestine and colon; CD49a in the prostate, mammary gland, and salivary gland; and many tissues showed mixtures or double-positive cells. This observation indicates that neither integrin is required for assignment to the core T_RM_ transcriptional program; instead, expression perhaps reflects tissue-specific adaptation. CD73^+^CD39^−^ expression helped distinguish CD8.Q from the adjacent exhaustion-associated clusters CD8.R and CD8.S (**Fig. 5f** and **Extended Data Fig. 4b**), which will be discussed in the following section. Thus, no single surface-marker combination was universally sensitive (fraction of true CD8.Q cells recovered) and specific (fraction of non-CD8.Q cells correctly excluded) for CD8.Q. The combination CD62L^−^CD44^+^CD69^+^CD73^+^CD39^−^ with variable CD103/CD49a provided the best practical compromise (median sensitivity 44%, improving to 73-78% positive predictive value when CD73^+^CD39^−^ was included; **Extended Data Fig. 4c,d** and **Extended Data Table 6** for comparison of different gating strategies). Flow cytometry validation on post LCMV-Armstrong P14 memory cells revealed tissue-to-tissue variation in CD73 and CD39 expression, while CD103/CD49a patterns largely validated the CITE-seq data (**Extended Data Fig. 4e**).

Key transcription factors showed expected patterns: *Hic1*^6,54^ and *Zfp683*^52,54^ (encoding Hobit) were enriched in cluster CD8.Q, while *Runx3*^53,54^ was broadly elevated across antigen-experienced clusters (**Extended Data Fig. 4f**), consistent with its conserved role in sustaining cytotoxic/residency programs^53,55,56^. Two smaller clusters, CD8.T (expressing *Hic1*) and CD8.wV (expressing both *Hic1* and *Zfp683*), shared parts of the T_RM_ transcriptional network and were enriched in intraepithelial lymphocyte (IEL)-like populations in gut and mammary gland (**Extended Data Fig. 4g,h**). Despite convergence on one core T_RM_ state, gene-program analysis within cluster CD8.Q from day 30 LCMV-Armstrong revealed tissue-driven heterogeneity. The T_RM_-like GP26 (e.g., *Cxcr6*, *Nkg7*, *S100a6*) was enriched in prostate and salivary gland, while the TCR-response GP35 (e.g., *Nr4a1*, *Klf6*, *Fos*, *Jun*) was selective for small intestine epithelium and lamina propria (**Fig. 5j**). These differences could reflect local microenvironmental cues such as TGF-β in the intestine^4,57^ or CXCL16 in salivary gland/prostate^58^.

In conclusion, antigen-specific T_RM_ cells across diverse non-lymphoid tissues converge on a single dominant transcriptional state (CD8.Q) in the immgenT framework, despite surface-marker and gene-program heterogeneity driven by tissue-specific adaptations. We provide a practical resource for interrogating CD8.Q tissue-specific marker expression across tissues through the Rosetta2 web tool (see **Methods**).

### Mapping exhaustion states in chronic antigen-driven CD8^+^ T cell responses

Clusters CD8.R and CD8.S were strongly enriched under conditions of persistent antigen exposure, including chronic LCMV-Clone 13 infection, multiple tumor models (B16 melanoma, KP lung adenocarcinoma, PDAC), and autoimmune settings (**Fig. 6a**, **Fig. 2c**, and **Extended Data Fig. 1h**). Notably, cluster CD8.Q, while primarily associated with tissue-resident memory after acute infection, also contains TIL, consistent with shared residency gene-expression programs between protective T_RM_ and some tumor-infiltrating populations^31,32,53,59–63^. Projection of independent TIL datasets (B16 melanoma and MC38) and T_EX_ cells from chronic LCMV-Clone 13 (ref. ^64^) using T-RBI confirmed localization primarily to clusters CD8.Q, CD8.R, and CD8.S (**Extended Data Fig. 5a,b**). CD8.Q additionally appeared in other persistent infection models (e.g., MNV-CR6; **Extended Data Fig. 5c**).

**Figure 6.**
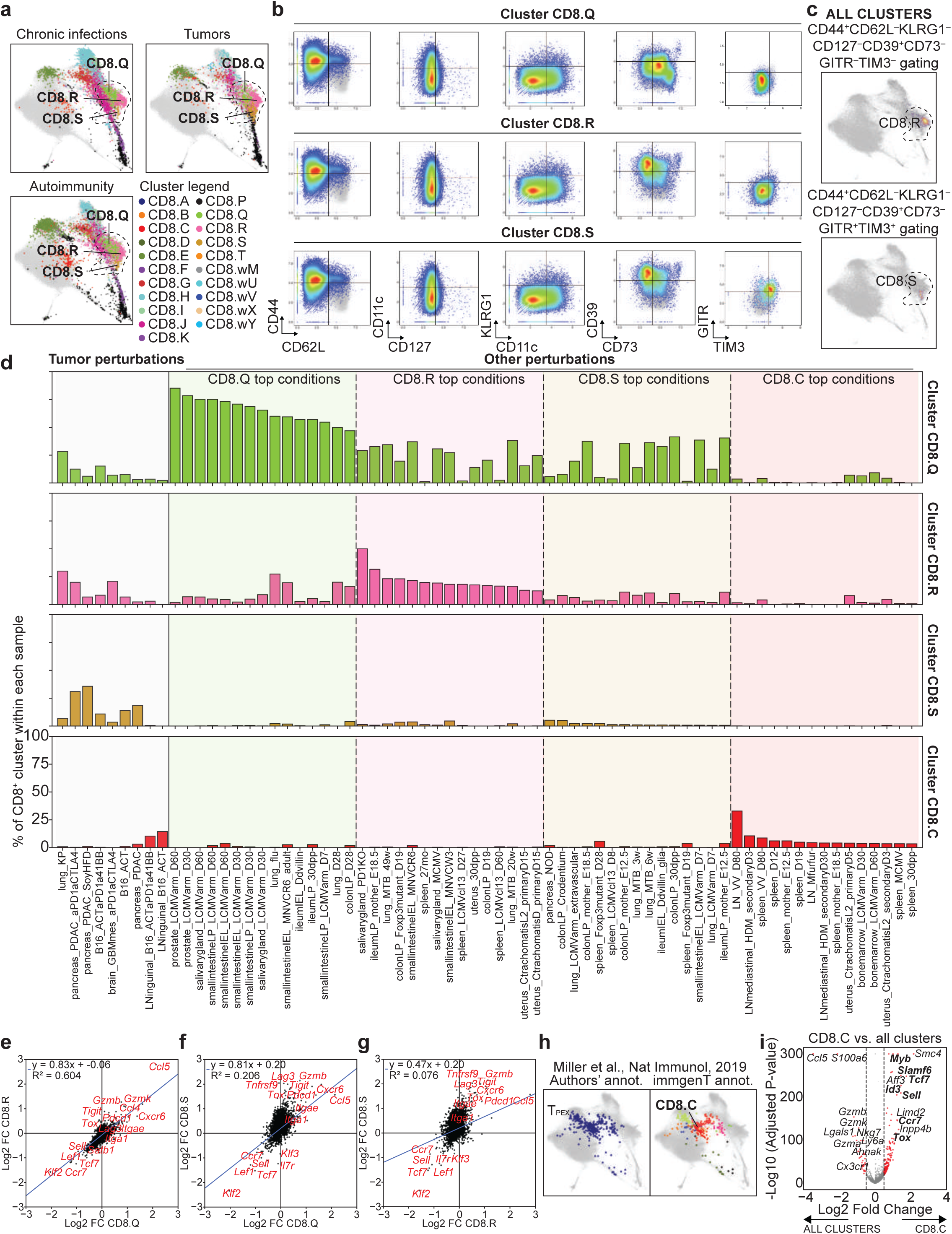
Exhaustion and residency-associated states form a continuum. **a**, CD8-MDE showing CD44^+^ clusters from chronic infection, tumor, and autoimmune conditions. Cells are colored by cluster identity; background immgenT-CD8 are shown in gray; **b**, Scatter plots showing predicted CITE-seq (log1p(CP10K)) surface-protein gating strategies for clusters CD8.Q, CD8.R, and CD8.S (key markers: CD44, CD62L, CD11c, CD127, KLRG1, CD73, CD39, GITR, TIM-3); **c**, CD8-MDE highlighting cells bioinformatically gated using the CD8.R (top) and CD8.S (bottom) gating strategies (shown above each plot); **d**, Bar plot showing the frequency of clusters CD8.Q, CD8.R, CD8.S, and CD8.C across top samples for tumors and other perturbations as proportion within each sample. Only samples with at least 10 total CD8^+^ T cells recovered in at least one cluster are shown; **e-g**, Fold-change versus fold-change plots comparing gene expression between clusters CD8.Q and CD8.R (**e**), CD8.Q and CD8.S (**f**), and CD8.R and CD8.S (**g**). Shared genes lie along the diagonal, whereas genes preferentially enriched in specific clusters fall along the horizontal or vertical axes, respectively. Selected genes are labeled; **h**, T-RBI (immgenT reference-based integration) of LCMV-specific cells in mice chronically infected with LCMV-Clone 13 on day 28 post infection from Miller et al.^64^. Only progenitor-exhausted (T_PEX_) cells, as annotated by the authors, are shown. Cells are colored by the authors’ annotation (left) and immgenT cluster identity (right); background immgenT-CD8 are shown in gray; **i**, Volcano plot showing the CD8.C-specific gene signature (versus all other clusters), highlighting T_PEX_-like features (*Tcf7*, *Tox*, *Sell*, *Ccr7*). **Abbreviations**: annot., annotation; vs., versus.

CITE-seq distinguished these three clusters from effector and memory CD8^+^ T cell states (**Fig. 6b**) by their lack of CD11c, KLRG1, and CD127 expression. In addition, four additional markers distinguished these clusters from one another: CD8.R was CD44^+^CD62L^−^CD11c^−^CD127^−^KLRG1^−^CD73^−^CD39^+^ (lacking GITR and TIM-3); CD8.S shared this profile but co-expressed GITR and TIM-3^65^; CD8.Q was CD73^+^CD39^−^ (lacking GITR and TIM-3). These combinatorial patterns enabled flow-cytometry-like gating that mapped back to the correct regions of the MDE (**Fig. 6c**).

The three clusters showed distinct contextual preferences (**Fig. 6d**). Cluster CD8.Q was abundant in non-tumor settings, especially after acute infection had resolved, but was also present in tumors. CD8.R appeared in multiple persistent-antigen infectious models (chronic LCMV, MCMV, MTB, MNV-CR6) and tumors. While CD8.S was almost exclusively enriched in tumor microenvironments and absent in chronic infections. Notably, exhaustion-like states (clusters CD8.R and CD8.S) remained relatively rare in most autoimmune models examined despite persistent self-antigen, raising the possibility that antigen persistence alone is insufficient to drive these states.

Transcriptionally, cluster CD8.Q was most similar to CD8.R (R^2^ = 0.604), while CD8.S was more divergent (R^2^ = 0.206) (**Fig. 6e-g**). Canonical exhaustion-associated genes (*Lag3*, *Tigit*, *Pdcd1*, *Tox*) were not exclusive to these clusters but showed progressively stronger differences from cluster CD8.Q to CD8.R to CD8.S (**Extended Data Fig. 5d**). These genes were detected in multiple exhaustion-like gene-programs (GP12, GP81, GP110), with CD8.S showing the highest activity (**Extended Data Fig. 5e,f**). Additionally, T_RM_-associated signatures^6,52,53^ were expressed at comparable levels in clusters CD8.Q, CD8.R, and CD8.S (**Extended Data Fig. 5g-i**), which underscores molecular convergence between residency and exhaustion programs operating in immunologically distinct contexts such as acute infection and tumor^31,32^. For example, direct comparison of cluster CD8.Q OT-I cells from flu-OVA lung (acute infection) versus KP lung tumor showed minimal transcriptional differences (**Extended Data Fig. 5j,k**), with similar surface-protein profiles (**Extended Data Fig. 5l**). Past work has indicated that TIL in the KP lung model are refractory to ICB but can be reinvigorated by cytokines^66,67^, emphasizing that understanding these differences will be important for developing new therapeutics. TCR clonal analysis revealed partial sharing among clusters CD8.Q, CD8.R, and CD8.S in B16 and KP tumors, suggesting common progenitors or dynamic intratumoral transitions (**Extended Data Fig. 5m**).

Mapping progenitor-exhausted (T_PEX_) cells provided a rigorous test of the immgenT data comprehensiveness. T-RBI projection of a published LCMV-Clone 13 T_PEX_ dataset^64^ mapped T_PEX_ primarily to cluster CD8.C, a rare cluster in individual immgenT samples but robust in the whole dataset with 5,000 total cells (**Fig. 6h**). Indeed, CD8.C displayed a T_PEX_-like profile (*Tcf7*, *Tox*, *Sell*, *Ccr7*) (**Fig. 6i**) and was enriched in B16 tumor-draining lymph nodes and, unexpectedly, in late (80 days post-infection) vaccinia virus memory lymph nodes (**Fig. 6d**).

In summary, clusters CD8.R and CD8.S represent distinct states that arise predominantly under persistent antigenic stimulation in chronic infection and cancer, while CD8.Q bridges T_RM_ and certain TIL populations. These states share core residency programs, yet their relative abundance varies across models. CITE-seq-guided combinatorial markers and T-RBI projection provide practical tools to annotate these states reproducibly between different datasets.

### immgenT as a comprehensive framework for mapping T cell data

The T-RBI analyses in **Fig. 4-6** and **Extended Data Fig. 1,2,5** illustrate and validate the T cell states that compose the immgenT-CD8 dataset. To illustrate the practical utility of the immgenT-CD8 framework, we tested its ability to annotate new datasets using T-RBI. We present four independent case studies spanning conditions both included and absent from the immgenT compendium. In the first case, we projected a published dataset of a hepatitis B virus (HBV) mouse model, which recapitulates key features of intrahepatic CD8^+^ T cell exhaustion observed in chronic HBV patients^68^ (**Fig. 7a**). Virtually all cells (>99%) mapped with high confidence to existing immgenT clusters, with no emergence of novel states (**Fig. 7b,c** and **Extended Data Fig. 6a,b**). For example, the authors’ T_EX_ population was captured predominantly by cluster CD8.R (89%), while cytotoxic cells mapped strongly to CD8.Q (82%) with memory cells enriched in cluster CD8.E (79%). These mappings (**Extended Data Table 7**) align with the expected distribution of exhaustion and residency programs under conditions of persistent hepatic antigen exposure and show how the immgenT-CD8 framework can be leveraged, through T-RBI, to quickly and precisely annotate CD8^+^ T cell states from unrelated conditions, allowing comparisons across unrelated datasets.

**Figure 7.**
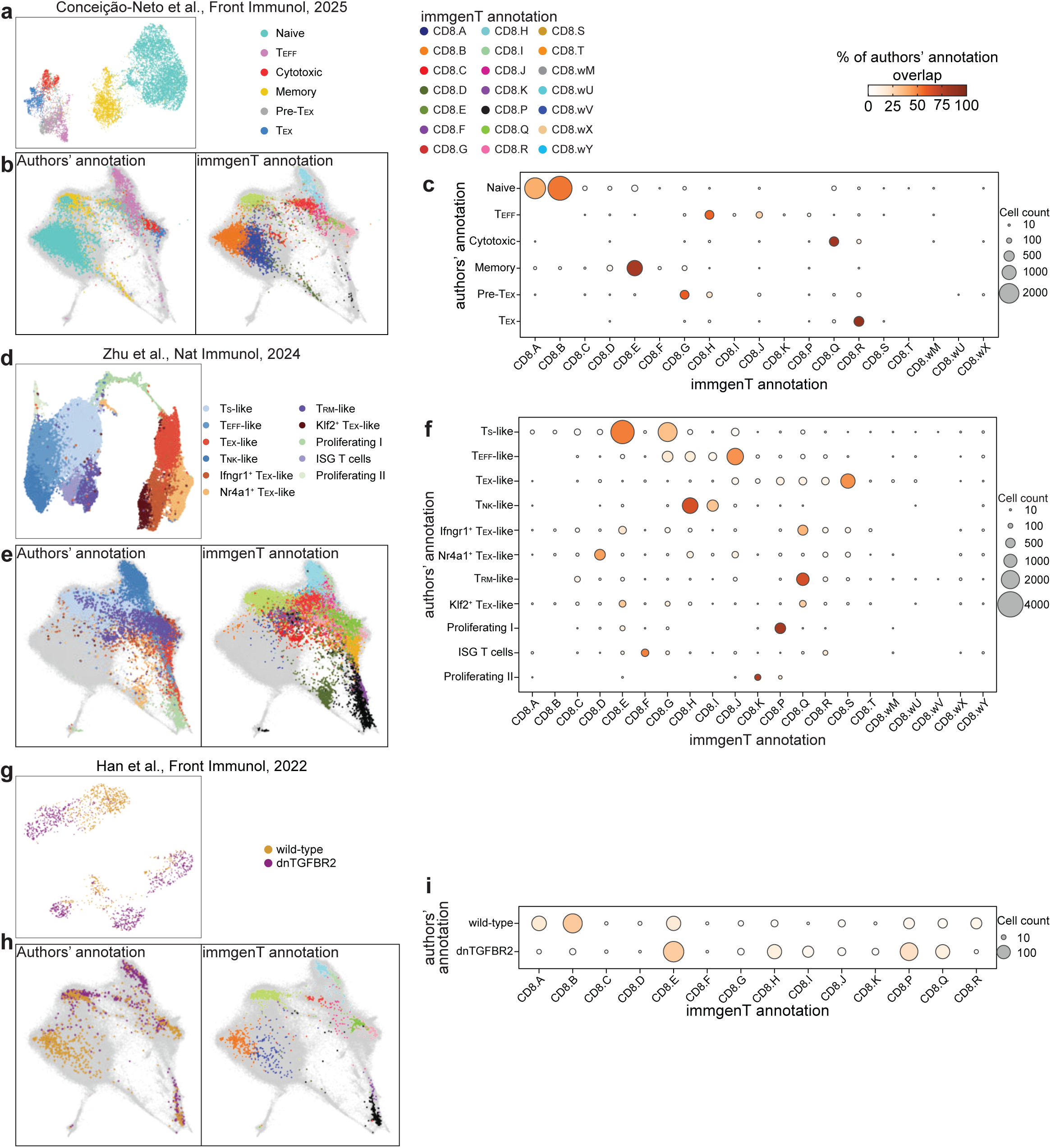
T-RBI analysis enables rapid, reproducible annotation of new CD8^+^ T cell datasets onto the immgenT reference. **a**, Original UMAP from Conceição-Neto et al. (2025)^68^ showing 6 annotated clusters of mouse intrahepatic CD8^+^ T cells from the AAV-HBV mouse model; **b**, The same CD8^+^ T cells projected onto the immgenT CD8-MDE (gray background) using T-RBI and colored by the authors’ original cluster annotations (left) or by immgenT cluster assignment (right); **c**, Dot plot showing the distribution of the authors’ original clusters across immgenT clusters (each dot represents the proportion of cells from one original annotation within an immgenT cluster; see **Extended Data Table 7**); **d**, Original UMAP from Zhu et al. (2024)^69^ showing 11 annotated clusters of mouse anti-CD19 CAR CD8^+^ T cells; **e**, The same CAR T cells projected onto the immgenT CD8-MDE (gray background) using T-RBI and colored by the authors’ original annotations (left) or by immgenT cluster assignment (right); **f**, Dot plot showing the distribution of the authors’ original clusters across immgenT clusters (see **Extended Data Table 8**); **g**, Original UMAP from Han et al. (2022)^70^ showing 2 annotated conditions of intrahepatic CD8^+^ T cells from dominant-negative TGFβRII transgenic mice that spontaneously develop primary biliary cholangitis versus wild-type; **h**, The same CD8^+^ T cells projected onto the immgenT CD8-MDE (gray background) using T-RBI and colored by the authors’ original cluster annotations (left) or by immgenT cluster assignment (right); **i**, Dot plot showing the distribution of the authors’ original clusters across immgenT clusters (each dot represents the proportion of cells from one original annotation within an immgenT cluster). **Abbreviations**: dn, dominant-negative.

In a second case, we projected a published dataset of mouse anti-CD19 CAR CD8^+^ T cells generated in the context of B-cell acute lymphoblastic leukemia (B-ALL) and CD19-expressing B16 melanoma^69^. The original study defined 11 clusters using standard unsupervised analysis (**Fig. 7d**). After T-RBI projection onto the immgenT reference, 96% of CAR T cells received high-confidence assignments to existing clusters (**Extended Data Fig. 6c**), with the small unassigned fraction (4%) being randomly distributed across the original dataset rather than originating from any specific cluster (**Extended Data Fig. 6d**). The projection did not identify novel states and preserved the internal diversity of the dataset while providing greater granularity and standardized labels (**Fig. 7e**). For example, the authors’ “stem-like” (TS-like) population split into two dominant immgenT states: 52% to cluster CD8.E (T_CM_-like) and 33% to CD8.G (T_EM_-like). “T_EX_-like” cells distributed across clusters CD8.Q, CD8.R, and CD8.S, as expected. “Effector-like” cells mapped mainly to CD8.J (47%) and actively proliferating effector-like cells (“Proliferating II”) to CD8.K (76%). “T_RM_-like” cells converged predominantly on cluster CD8.Q (66%) (**Fig. 7f** and **Extended Data Table 8**). This analysis demonstrates how the framework harmonizes annotations and reveals finer structure within previously defined populations.

In a third case, we evaluated the framework using a published dataset of hepatic CD8^+^ T cells isolated from the dominant-negative TGFβRII transgenic mouse model of primary biliary cholangitis and wild-type controls^70^ (**Fig. 7g**). CD8^+^ T cells from transgenic mice were prominently distributed across clusters CD8.E (29%), CD8.P (22%), CD8.H (14%), and CD8.Q (13%), consistent with increased effector, proliferating, and tissue-resident/exhaustion-like states in the diseased liver (**Fig. 7h,i**). In contrast, wild-type cells showed greater enrichment in the resting clusters CD8.B (30%) and CD8.A (17%). T-RBI assigned the majority of cells (98.5%) with high confidence to the existing 21 clusters (**Extended Data Fig. 6e,f**). Together, these results illustrate that the immgenT reference can resolve disease-associated remodeling of the CD8^+^ T cell landscape while providing standardized, biologically interpretable annotations.

In the final case, we projected the scRNA-seq layer of a large paired scRNA/scATAC-seq dataset^71^ of TCR-transgenic P14 CD8^+^ T cells across LCMV-Armstrong, LCMV-Clone 13, and four GP33-expressing tumor models (**Extended Data Fig. 6g**). While LCMV-Armstrong, LCMV-Clone 13, KP, and B16 models are represented in immgenT, this study also includes B-ALL and triple negative breast cancer tumor models that are absent from immgenT. Nearly all cells (99%) mapped with high confidence to existing immgenT clusters (**Extended Data Fig. 6h,i**), with no emergence of novel states, further supporting the comprehensiveness of the reference. T-RBI again improved granularity: the authors’ “T_CM_” population split between clusters CD8.E (38%) and CD8.G (43%), while their “T_R_-tem” and “T_RM_” populations both converged strongly on cluster CD8.Q (**Extended Data Fig. 6j,k** and **Extended Data Table 9**). These examples show that T-RBI can both refine existing annotations and converge disparate labels onto shared molecular states.

To facilitate broader use, we developed a practical 13-marker CITE-seq-compatible flow cytometry panel predicted to sample all 21 clusters (**Extended Data Fig. 7a,b**). Most markers showed strong RNA/protein concordance, although notable discordance was observed for CD44 and CD69, reinforcing the value of combined measurements and the need for careful validation of flow cytometry reagents when assigning states (**Extended Data Fig. 7c**). We also defined and flow-validated 21 cluster– or family-specific gating strategies (**Fig. 3e** and **Extended Data Fig. 2b-e, 3a, 4e, 7d,e**), reporting sensitivity, specificity, positive predictive value, and negative predictive value for each (**Extended Data Fig. 7f** and **Extended Data Table 4**). A minimal flow cytometry panel with cluster– or family-specific markers, suggested fluorochromes, and specific recommendations is provided in **Extended Data Table 10**. An interactive web tool, Rosetta2 on the immgenT website, allows users to perform in silico gating on the full 128-marker CITE-seq panel and visualize the gated cells directly on the CD8-MDE. Together, these resources (including T-RBI projection, immgenT-CD8 flow cytometry panel, Rosetta2, and more as described in the **Methods**) enable rapid, reproducible annotation of new mouse CD8^+^ T cell datasets onto the unified immgenT coordinate system.

## DISCUSSION

The immgenT-CD8 framework provides a comprehensive, molecular reference and single-cell-resolution coordinate system for mouse CD8^+^ T cell differentiation encompassing CD8^+^ T cells across a wide breadth of immunological contexts. Twenty-one shared transcriptional and proteomic states recurred across 53 immune perturbations, 45 tissues, and remained robust in external datasets. Classical models (acute LCMV-Armstrong for effector-to-memory transitions, chronic LCMV-Clone 13 or tumor challenge for exhaustion-like states) mapped to the atlas with the same states generalizing remarkably well across diverse viral, bacterial, parasitic, and fungal infections, tumor, autoimmunity, and aging. Although variable representations of nearly all 21 clusters exist at baseline, their proportions are dramatically reshaped by immunological challenge, indicating that CD8^+^ T cell states are context-dependent rather than fixed cell types. These findings reveal greater molecular convergence than classical subset definitions imply, with effector and memory programs unexpectedly reused across both protective and dysfunctional settings.

CD8^+^ T cell differentiation across the immgenT landscape was largely dictated by two major axes: acute versus chronic antigen exposure and lymphoid versus non-lymphoid residence, with time and tissue microenvironment imposing additional fine-tuning. Acute resolving challenges generate transient effectors (clusters CD8.I/J/K) that transition into circulating memory (clusters CD8.E-H) or tissue-resident memory CD8^+^ T cells (CD8.Q). Persistent antigen in chronic infection or malignancy drives progenitor and terminally-exhausted compartments (clusters CD8.C, and CD8.R/S, respectively). Tissue context exerts particularly strong influence on cluster CD8.Q (canonical T_RM_), where surface integrin usage (CD103 versus CD49a) and gene-program use vary across organs yet converge on a unified transcriptional identity. Circulating T cell states exhibit tighter coherence, underscoring tissue-imposed reprogramming as a likely source of CD8^+^ T cell heterogeneity.

A striking observation is the low frequency of canonical exhaustion-like states (clusters CD8.R and CD8.S) in autoimmune models despite chronic inflammation and persistent self-antigen. Although exhaustion-like profiles have been described for some autoreactive T cells, potentially restraining aberrant responses^72–75^, this suggests differences in antigen perception and response in autoimmunity versus chronic viral infection or tumor, possibly due to deletional tolerance, altered presentation, insufficient co-stimulation, or active suppression as additional mediating factors. T-RBI offers an immediate tool to interrogate additional autoimmune datasets for cryptic exhaustion-like populations. Another key consideration when interpreting exhaustion-associated states (CD8.Q/R/S) is the substantial variability in antigen load, duration, and tissue context across the models included in the dataset. For example, chronic LCMV-Clone 13, different transplantable tumors (B16, KP, PDAC), and autoimmune settings differ markedly in antigen persistence and inflammatory milieu. This variability may contribute to the observed differences in abundance and dominance of clusters and underscores why direct functional equivalence cannot be assumed across models.

The global MDE highlights both the power and sensitivity of the framework: it appears continuum-like when superimposing hundreds of conditions, yet discrete clusters remain robust in RNA, protein, and sample space. Cluster-defining signatures were conserved across vastly different contexts, capturing convergent molecular endpoints rather than idiosyncrasies. Cluster CD8.Q exemplifies this observation, reconciling extensive tissue-driven heterogeneity within a single coherent state. Immunologists have long sought unequivocal cell-surface markers; this complex, comprehensive dataset explains the elusiveness of a single marker as most CD8^+^ T cell states require combinatorial signatures and coordinated gene-programs rather than single proteins (see also immgenT-Cosmology^34^ for additional discussion on the interpretation of the clusters).

While alternative approaches (different integration methods, higher resolution clustering, or condition-specific re-analysis) could potentially reveal additional nuance or condition-dependent substructure, our saturation analysis, cross-dataset projections, and biological coherence across all the immunological challenges argue that the major recurrent states are robustly captured in the current immgenT framework. We therefore propose this reference as a stable coordinate system for harmonizing CD8^+^ T cell states and annotations across the field.

Caution is warranted in assigning rigid functional labels. Bona fide T_RM_ from acute infections map to cluster CD8.Q, yet subsets of TIL in melanoma and other chronic conditions occupy the same state, blurring the definitions of protective residency and dysfunctional persistence. Recent work shows tumor-resident exhausted T cells are ontologically, clonally, and functionally distinct from canonical T_RM_ despite shared residency features, with T_RM_ rewired to exhaustion under chronic antigen and T_EX_ unable to form conventional T_RM_ upon antigen withdrawal^31,32^. Similarly, clusters CD8.E-H recapitulate T_CM_, T_EM_, and t-T_EM_/LLEC after acute infection but identical states appear in tumor-draining LNs, autoimmune lesions, aged mice, and unchallenged animals. These findings argue against overly prescriptive nomenclature and favor context-aware, framework-based descriptions that acknowledge molecular convergence. The convergent nature of CD8^+^ states, reaching similar molecular endpoints via diverse trajectories dictated by context, suggests shared regulatory pathways amenable to therapeutic targeting across diseases.

Recent community guidelines for T cell nomenclature recognize the limitations of rigid subsets and advocate for a modular paradigm that denotes individual properties (e.g., stemness, residency, exhaustion) using concise descriptors^33^. The intent aligns with the goals of immgenT: the 21 clusters provide a molecular basis to anchor immune functions and context-specific interpretations. In immgenT, we leveraged molecular profiling from experiments where specific T cell functions have been robustly demonstrated (e.g., antigen-specific CD8^+^ T cells in the gut post LCMV infection shown to be resident via parabiosis) to map classical nomenclature (e.g., T_CM_, T_EM_, T_RM_, T_PEX_, T_EX_, etc.) onto the atlas. To support adoption of the reference, we provide a mapping between commonly used terms and the 21 immgenT states in **Fig. 4g** and **Extended Data Table 5**. The public resources (T-RBI, Rosetta2) extend utility: T-RBI harmonizes annotations and discovers rare subsets (e.g., placing T_PEX_ in cluster CD8.C and revealing its enrichment at late vaccinia memory). Rosetta2 enables in silico gating on the 128-marker CITE-seq panel. A streamlined 13-marker flow panel and validated strategies for all 21 states further democratize access (https://www.immgen.org/ImmGenT/).

Despite its breadth, the framework primarily maps “where” cells reside in molecular space rather than “how” they arrive there. Key open questions include transcription-factor cascades, epigenetic landscapes, and lineage relationships. Integrating trajectory inference, CRISPR screens, ATAC-seq, and fate-mapping with T-RBI will transform this static map into a dynamic understanding of CD8^+^ T cell differentiation.

In summary, the immgenT-CD8 framework provides an approach to resolve decades of disparate literature into a cohesive reference. By providing robust states, validated markers, and publicly accessible tools, it equips the field to annotate heterogeneity reproducibly, uncover unexpected biology, and accelerate translational efforts in vaccination, tumor immunotherapy, and chronic infection.

## METHODS

### Mice

Mice used in the immgenT dataset are described in detail in the immgenT-Cosmology manuscript^34^ (**Extended Data Table 1**) and on the immgenT website, including sex, age, and genetic background. With rare exceptions, experiments were performed using C57BL/6 (B6) mice, most of which were sourced from The Jackson Laboratory. Experimental conditions, including infection models, immunization strategies, and tissue processing protocols, are detailed in the immgenT-Cosmology manuscript (**Extended Data Table 1**), the GEO GSE297097 dataset, and the immgenT website. Both male and female mice were used.

MC38-SIY and B16-SIY tumor experiments were performed using female C57BL/6 mice (6-8 weeks old) obtained from Taconic Biosciences and housed under SPF conditions at the MIT Koch Institute animal facility. MC38-SIY colon carcinoma and B16-SIY melanoma cell lines (Gajewski Laboratory, University of Chicago) were cultured in DMEM supplemented with 10% FBS, 1% penicillin/streptomycin, and 1× HEPES at 37°C, 5% CO_2_. Cell lines were routinely tested for mycoplasma.

The flow cytometry experiments presented in the present study were conducted on mice bred and maintained under specific pathogen-free conditions at the University of California, San Diego, in accordance with procedures reviewed and approved by the University of California, San Diego Institutional Animal Care and Use Committee (IACUC) guidelines (Protocol ID: S04105).

### immgenT dataset: experiments and data processing

The immgenT dataset comprises 66 experiments of single-cell RNA-seq, CITE-seq (128-plex), and paired TCRαβ sequencing (10x Genomics 5′ v2 platform), corresponding to 80 encapsulation runs. Each encapsulation is assigned a unique “IGTx” identifier (IGT1-96) used to track datasets. In a minority of cases, two IGT identifiers denote parallel encapsulations of the same biological samples. Individual cells are indexed by a unique IGT.cellID, and samples are tracked using hashtag identifiers (IGT.HT). A detailed description of data acquisition and processing is provided in the immgenT-Cosmology manuscript^34^. Briefly:

#### Logistics

Experiments were conducted across multiple laboratories in the United States. Participating laboratories carried out mouse treatments (e.g., infection or immunization) and tissue processing with their established materials and reagents. The immgenT research assistant (Ian Magill) traveled to the site, helped with sample CITE-seq labeling and cell sorting, and performed encapsulation and library preparation. Each experiment typically included ∼10 hashtagged and pooled samples, and a standardized spleen control for batch-effect assessment. Library construction and sequencing were centralized at the Broad Institute.

#### Sample processing, library construction, and sequencing

Flow cytometry–sorted T cells were multiplexed using TotalSeq-C hashtag^76^ antibodies (BioLegend), enabling pooling prior to encapsulation. Cells were stained with a custom 128-antibody TotalSeq-C panel and subjected to joint single-cell RNA, surface protein (CITE-seq), and TCR sequencing using the 10x Genomics 5′ v2 platform. Libraries were sequenced on an Illumina NovaSeq.

#### Data processing and quality controls

Gene expression, protein, hashtag, and TCR count matrices were generated using Cell Ranger, and hashtag-based demultiplexing was performed in Seurat^77,78^. Cells were filtered based on RNA and protein quality-control metrics detailed in the immgenT-Cosmology manuscript (Extended Data Note 1).

#### Data integration

Datasets were integrated using totalVI^79,80^, with the 10x Genomics lane (IGT) specified as a batch covariate. The model was trained on all detected genes and proteins using a 30-dimensional latent space (default parameters).

#### Dimensionality reduction

Dimensionality reduction was performed using Minimum Distortion Embedding (MDE) implemented in the pyMDE^81^ library via the pymde.preserve_neighbors() function with default parameters. In contrast to UMAP, which is stochastic and graph-based, pyMDE preserves both local neighborhood structure and global geometry with minimal distortion. After computing the embedding on the full immgenT dataset (IGT1-96), coordinates were reused to anchor immgenT cells as a reference, enabling projection of new datasets without altering the original embedding. This enabled the construction of shared T cell reference embeddings (both all-T and lineage-specific, including CD8-specific embeddings).

#### Cell clustering and annotation

Clustering was performed in the totalVI latent space using the Louvain algorithm (Seurat FindClusters). Clustering solutions were evaluated across resolutions (0.5-4) and optimized using silhouette scores to balance over– and under-clustering. Clusters were merged or split based on consistency across samples and coherence of RNA and protein expression profiles (see immgenT-Cosmology manuscript, Extended Data Note 3). In total, 1,710 of 719,580 cells (0.02%) could not be confidently annotated and were labeled as “unclear” and excluded. Small provisional clusters (<1% of cells) are denoted with a “w” prefix. CD8αβ T cells were identified as CD8α^+^CD8β^+^ T cells based on combined RNA and protein expression of canonical markers and transcription factors, including Cd3e, Trbc1, Trbc2, Cd8a, Cd8b, as well as surface proteins CD3, TCRβ, CD8A, and CD8B. For CD8αβ cells, we generated a CD8^+^-focused MDE and performed unbiased clustering that yielded 21 recurrent states after saturation was reached (approximately half the experiments).

#### CITE-seq analysis

The full panel of 128 antibodies and associated quality metrics are provided in Extended Data Table 4 and Extended Data Note 1 of the immgenT-Cosmology manuscript^34^. Antibody performance was evaluated using RNA-protein correlation and protein dynamic range, stratifying antibodies into high-, intermediate-, and low-performance groups (n = 61, 41, and 26, respectively). Overall, most of the CITE-seq panel performed robustly, enabling dropout-resistant protein quantification that closely parallels flow cytometry and supports both high-dimensional analyses and conventional gating strategies. For flow-cytometry-like plots, protein counts were normalized using Seurat’s LogNormalize method (log1p CP10K).

#### UMAP and clustering of individual IGT datasets

UMAP and clustering were also performed on individual IGT datasets, generating dataset-specific UMAPs used in the Rosetta2 databrowser and in some immgenT manuscripts. These used the standard Seurat functions: NormalizeData(normalization.method = “LogNormalize”, scale.factor = 1e4) %>% FindVariableFeatures(selection.method = “vst”, nfeatures = 2000) %>% ScaleData(features = VariableFeatures(.)) %>% RunPCA(features = VariableFeatures(.), npcs = 50) %>% FindNeighbors(dims = 1:30) %>% FindClusters(resolution = 1).

### immgenT Reference-Based Integration (T-RBI)

The full methods are described in the immgenT-Cosmology manuscript^34^. Integration of query datasets using T-RBI is available at https://www.immgen.org/ImmGenT/.

Reference-based integration of external datasets (T-RBI) was performed using scVI/scANVI^80,82,83^, and pyMDE. Query datasets are first filtered for T cells using gene signature scoring, with γδ T cells identified and analyzed separately from αβ T cells. scVI models were trained jointly on reference (immgenT) and query data to learn a shared latent space, followed by scANVI to predict lineage and sub-lineage annotations with associated confidence scores. Cells with low-confidence predictions were iteratively reassigned until convergence. To quantify whether unannotated cells occupy regions of the embedding space not represented in the annotated reference (i.e., potential novel T cell states), we defined a discovery score based on a local k-nearest neighbor (kNN) distance ratio between query and reference cells. Scores >1 indicate that a cell is locally closer to other unannotated cells than to annotated cells, consistent with occupancy of under-represented or novel regions of the reference space, whereas scores ≤1 indicate embedding within previously annotated regions. The resulting latent representations were then used to project query cells onto the immgenT reference embeddings using pyMDE with anchored constraints, preserving the original atlas structure while incorporating new data.

Final outputs include lineage annotations, confidence scores, discovery scores, and coordinates in both global (allT-MDE) and CD8-specific MDE embeddings, enabling direct comparison of query datasets within the immgenT framework.

### Differential gene expression

#### Limma

Differential gene expression was performed using a pseudobulk linear modeling framework based on limma^84,85^ and edgeR. Briefly, raw RNA counts from the Seurat object were aggregated by cell cluster (annotation_level2) and experiment (IGTHT), and normalized using trimmed mean of M-values (TMM). A design matrix encoding cluster–experiment combinations was constructed, and gene-wise linear models were fit to the normalized expression matrix using limma. Differential expression was assessed using empirical Bayes moderation of variance estimates, and statistical significance was determined with Benjamini–Hochberg correction for multiple testing (limma_wrapper_template.sh; limma_make_tmm_template.R; limma_fit_template.R; limma_contrasts_template.R).

#### FlashierDGE

Fast differential expression was performed in the flashier^86,87^ EBMF semi-NMF framework by leveraging the learned gene-programs (factors) and cell loadings from a 200-factor model fit to log-normalized expression. For a given comparison, mean factor loadings were computed separately for group 1 and group 2, and the difference in mean loadings (Δloadings) provided differential gene-program activity. Gene-level differential expression was then reconstructed directly from the factorization as F×Δloadings, yielding per-gene log fold changes. The same framework also returned average expression estimates for both programs and genes from the group-wise mean loadings and reconstructed group means. The function is available in ZemmourLib package (FlashierDGE).

### TCR clonotype sharing and overlap analysis

Clonotype sharing across CD8^+^ T cell clusters was assessed using matched single-cell TCR sequencing data as described in the immgenT-TCR manuscript^88^. Clonotypes were defined based on paired TCRα and TCRβ sequences with full nucleotide-level identity in the junction sequences. For each cluster, the proportion of cells with expanded clonotypes was calculated as the fraction of cells with clonotypes observed more than once in the sample. To quantify overlap between clusters, pairwise clonotype sharing was computed using the Jaccard index on sets of unique clonotypes per cluster. For visualization, a weighted network of cluster relationships was constructed, where edge weights correspond to the Jaccard index. Edges corresponding to only one shared clonotype were omitted for clarity and the network was overlaid onto the MDE embedding using cluster centroids.

### Gene-programs and empirical Bayes matrix factorization of gene expression

To identify latent transcriptional programs, we applied empirical Bayes matrix factorization (EBMF) with a semi-non-negative matrix factorization (semi-NMF) structure to log-normalized single-cell RNA-seq expression using the flashier framework^86,87^. Briefly, EBMF decomposes an expression matrix into a sum of low-rank components while learning sparsity, shrinkage, and noise parameters directly from the data via empirical Bayes, enabling robust identification of interpretable gene-programs without requiring a priori selection of factor number or regularization strength. In the semi-NMF formulation used here, gene loadings are constrained to be non-negative, facilitating biological interpretability of gene-programs, while cell loadings are unconstrained.

Gene expression was log-normalized using Seurat’s LogNormalize() with a scale factor set to the mean total UMI count per cell (mean(nCount_RNA)), yielding log1p-transformed expression values. Genes were filtered prior to model fitting to reduce technical and clonotype-driven variation: TCR variable/constant genes (e.g., Trbv/Trbj/Trbc, Trav/Traj/Trac, Trgv/Trgc, Trdv/Trdc), mitochondrial genes (^mt-), ribosomal genes (including Rpl/Rps/Mrpl/Mrps/Rsl), predicted/RIKEN/pseudogene-like features (including Gm genes, genes ending in Rik, and –ps pseudogenes), and genes not detected in any cell (row sum of counts = 0) were excluded.

Variance regularization parameters were set using a reference Poisson sampling procedure: a Poisson distribution with rate 1/n (where n is the number of cells) was sampled to estimate the standard deviation of log(x+1) and used as the residual noise scale parameter (S). EBMF components were fit using flash() (flashier^86,87^) with a mixture of empirical Bayes priors (ebnm_point_exponential and ebnm_point_laplace) and variance model var_type = 2, with a maximum of 200 greedily added factors (greedy_Kmax = 200). Backfitting was enabled to refine factor estimates.

### immgenT-CD8 flow cytometry panel design and testing

#### Panel design

For each lineage, COMETSC^89^ was applied to normalized protein expression matrices together with MDE coordinates and cluster annotations, allowing up to two-gene combinations (-K = 2) to identify combinatorial markers. In combination with literature curation, extended marker combinations were evaluated for CD8^+^ cluster specificity using CITE-seq by mapping onto the CD8-MDE and testing candidate strategies by flow cytometry.

#### Single-cell suspensions

To identify CD8^+^ T cells in the vasculature of non-lymphoid tissues (small intestine, lung, and salivary gland), 3 µg of anti-CD8α (clone 53-6.7, Thermo Fisher Scientific Cat. #47-0081-82) conjugated with APC-eFluor780 were injected i.v. into the mice 3 minutes before sacrifice. Cells that displayed weak or no labeling with the anti-CD8α antibody (i.v.^−^) were considered to be located outside the vasculature. Single-cell suspensions of spleens were prepared by mechanical disruption, followed by ACK Lysing Buffer for 5 minutes at room temperature. For tissue preparations, small intestine was prepared by excision of Peyer’s patches and removal of luminal contents. To collect the intraepithelial lymphocytes (IEL) the entire tissue section was cut longitudinally, then cut laterally into 1 cm pieces, which were then incubated at 37°C for 30 minutes in Hanks’ balanced salt solution with 2.4 mg/ml of Hepes, 2.1 mg/ml of sodium bicarbonate, 8% bovine growth serum and 0.154 mg/ml of DTE (233152-5GM; EMD Millipore). The lung and salivary gland were cut into small pieces then incubated shaking at 37°C for 1 hour or for 30 minutes, respectively, in RPMI with 1.2 mg/ml of Hepes, 292 µg/ml of L-glutamine, 1 mM MgCl_2_, 1 mM CaCl_2_, 5% FBS and 100 U/ml of collagenase (LS004196; Worthington). Lymphocytes from non-lymphoid tissues were further enriched using a 44% and 67% Percoll (P1644-1L; Sigma) density gradient.

#### Staining

Up to 1 × 10^6^ cells were stained. Cells were first incubated with Live/Dead Blue (Thermo Fisher Scientific Cat. #L34962) for 20 min at 4 °C protected from light. After washing in FACS buffer (PBS, 2% bovine growth serum, 0.01% sodium azide), cells were incubated with Fc Block for 10 min, followed by surface antibody staining for 20 min at 4 °C in FACS buffer against CD45.1 (clone A20, BUV395, Thermo Fisher Scientific Cat. #363-0453-82), CD3 (clone 145-2C11, BUV496, BD Biosciences Cat. #612955), CD4 (clone RM4-5, BUV615, BD Biosciences Cat. #751486), CD69 (clone H1.2F3, BUV737, Thermo Fisher Scientific Cat. #367-0691-82), CD8β (clone YTS156.7.7, BUV805, BD Biosciences Cat. #755247), CD103 (clone 2E7, BV480, Thermo Fisher Scientific Cat. #414-1031-82), GITR (clone DTA-1, Super Bright 600, Thermo Fisher Scientific Cat. #63-5874-82), CD44 (clone IM7, BV650, Thermo Fisher Scientific Cat. #416-0441-82), KLRG1 (clone 2F1, Super Bright 702, Thermo Fisher Scientific Cat. #67-5893-82), TCRβ (clone H57-597, Super Bright 780, Thermo Fisher Scientific Cat. #78-5961-82), CD11c (clone N418, BB515, BD Biosciences Cat. #565586), CD49a (clone Ha31/8, PE, BD Biosciences Cat. #562115), CD39 (clone Duha59, PE/Dazzle594, BioLegend Cat. #143812), CD127 (clone A7R34, PE-Cy5, Thermo Fisher Scientific Cat. #15-1271-82), TIM-3 (clone RMT3-23, PE-Cy7, BioLegend Cat. #119716), CD73 (clone TY/11.8, APC, BD Biosciences Cat. #567499), CD62L (clone MEL-14, APC-R700, BD Biosciences Cat. #565159) (**Extended Data Table 10**). Brilliant Stain Buffer (Thermo Fisher, Cat. #00-440-942) was used in the staining solution when more than two BV-conjugated antibodies were included. All reagents were titrated prior to use to determine optimal concentrations.

#### Data acquisition and analysis

Data were acquired on a Cytek Aurora (Cytek Biosciences), BD LSRFortessa X-20 (BD Biosciences) or equivalent and analyzed with FlowJo v10 (TreeStar, BD LifeSciences).

### immgenT resource: public data browsers

Beyond the data and analyses reported in this and accompanying articles, immgenT’s goal as a public resource is served by a set of data browsers which enable multi-faceted exploration of the data. They are accessible through the immgenT portal (https://www.immgen.org/ImmGenT/) and include:

**<u>Rosetta2</u>** is an interactive single-cell viewer that connects RNA-seq and protein (CITE-seq) data. Cell populations can be visualized on parallel RNA and protein UMAPs, coloring reciprocally RNA– or protein-based clusters. The expression of individual genes, or of user defined gene signatures can be mapped on these representations. A cell gating interface modeled after flow cytometry serially generates 2D CITE-seq plots, on which cells can be “gated”, for projection onto UMAPs or computation of differential expression. Rosetta2 returns differentially expressed genes and proteins (as heatmaps or tables). Rosetta2 can be used to explore individual datasets or integrated cells grouped by lineage.

**<u>TCR Browser</u>** is a searchable interface for TCR sequences across the immgenT data. TCR repertoires can be browsed within immgenT datasets, searched for TCRs that use specific V, J or CDR3 elements, or for similarity to a user provided TCR.

**<u>T-RBI portal</u>** (Reference-Based Integration) allows AI-driven mapping of external single-cell RNAseq datasets into the immgenT framework. Users upload their own data and metadata files (standard sc formats), which are then mapped and annotated within the immgenT framework. After an overnight compute, the tool returns annotation tables with integration statistics and annotation of every cell in the immgenT cluster framework, and x/y coordinates in the reference MDE space.

**Skyline browser** represents, as a classic bar histogram, a pseudo-bulked overview of expression profiles of a given gene across all or selected immgenT cell clusters.

### Plotting

Plots were generated in R using ggplot2^90^, S-Plus, Seurat^78^, or the ZemmourLib R package (https://github.com/dzemmour/ZemmourLib, v0.1.3). Heatmaps were generated using pheatmap (v1.0.13) or Morpheus (Broad Institute)

### Code availability

Code is available at the following repositories: https://github.com/immgen/immgen_t_git/, https://github.com/dzemmour/immgent_rbi, https://github.com/dzemmour/immgen_t, and https://github.com/immgen/immgenT_Project. All other analyses were performed with standard bioinformatics pipelines and packages.

### Data availability and resources

All raw and processed sequencing data generated in this study are available through the Gene Expression Omnibus (GEO) under accession GSE297097 and Zenodo under DOI 10.5281/zenodo.21839962. MC38-SIY and B16-SIY tumor experiment data are available with GSE316401. Studies mapped using T-RBI: GSE244487, GSE122712, GSE131535, GSE164978, GSE202543, GSE186164, GSE129030, GSE109774 (TabulaMuris), GSE202543, GSE267584, GSE182275, GSE186333. Additional resources are described in detail in the immgenT-Cosmology manuscript (Extended Data Note 2). Gene signatures for each cluster of the CD8-MDE are listed in the **Extended Data Table 11** and the signature genes part of the gene-programs included in the immgenT-CD8 manuscript are listed in the **Extended Data Table 12**.

The immgenT portal (https://www.immgen.org/ImmGenT) provides lineage and cluster annotations (tissue distribution, sample enrichment, and gene signatures), and access to several analytical tools for the immgenT dataset:

– Pseudobulk gene expression across clusters can be explored using the immgenT Skyline;
– Individual experiments, as well as integrated datasets, can be interactively visualized using the Rosetta2 platform, which displays UMAP and MDE embeddings as well as flow-like scatter plots, gene expression, surface protein abundance, and differential expression analyses;
– TCR data from immgenT (https://rstats.immgen.org/tcrbrowser/);
– Mapping of external datasets onto the immgenT framework using T-RBI.

## ACKNOWLEDGEMENTS

We acknowledge the Flow Cytometry Core Facility and the Sequencing Core Facility at the La Jolla Institute for Immunology, and the Brown University Flow Cytometry Core for assistance with cell sorting. We thank the NIH Tetramer Core Facility (NIH Contract 75N93020D00005 and RRID:SCR_026557) for providing LCMV gp33-41 (KAVYNFATC) tetramer. National Institutes of Health grants: R24-072073 (ImmGen consortium), R01AI179952 (A.W.G.), R37AI067545 (A.W.G.), R01AI072117 (A.W.G.), R01AI150282 (A.W.G.), R01AI192333 (S.M.B.), R01AI172905 (S.M.B.), K00CA222711 (N.E.S.), F31AI176705 (K.K.T.), and F31DE032593 (S.M.B.). G.G. was a Cancer Research Institute Irvington Fellow supported by the Cancer Research Institute (CRI4145). A.M.G. was supported by a NOMIS Foundation Postdoctoral Fellowship. T.A.H. is supported by a postdoctoral fellowship from the Ludwig Center at MIT’s Koch Institute.

## AUTHOR CONTRIBUTIONS

Conceptualization: D.Z. Methodology: G.G., A.M.G., O.B., T.A.H., S.L., S.M.B., O.C., A.M., D.P., N.E.S., S.Q., K.K.T., A.F., K.P.C., E.D., T.S., D.Z. Investigation: G.G., A.M.G., O.B., T.A.H., S.L., S.M.B., O.C., A.M., D.P., N.E.S., E.D., T.S., S.S., S.M.B., S.M.K., A.W.G., D.Z. Visualization: G.G., D.Z. Funding acquisition: S.S., S.M.B., S.M.K., A.W.G., D.Z. Project administration: A.W.G., D.Z. Supervision: G.G., A.W.G., D.Z. Writing – original draft: G.G., A.W.G., D.Z. All the authors reviewed and edited the manuscript.

## COMPETING INTERESTS

Authors declare no competing interests.

## COLLABORATORS

### Participants in the immgenT Project include

Aaron Liu^1^, Alexander Chervonsky^2^, Alexandra Cassano^2^, Alia Welsh^3^, Amir Ferry^11^, Ananda Goldrath^11^, Andrea Lebron-Figueroa^5^, Ankit Malik^2^, Anna-Maria Globig^4^, Antoine Freuchet^2^, Bana Jabri^2^, Charlotte Imianowski^6^, Christophe Benoist^5^, Claire Thefaine^7^, Dan Kaplan^6^, Dania Mallah^5^, Dario Vignali^6^, David Sinclair^5^, David Zemmour^2^, Derek Bangs^8^, Domenic Abbondanza^2^, Enxhi Ferraj^9^, Eric Weiss^6^, Erin Lucas^7^, Evelyn Chang^9^, Gavyn Chern Wei Bee^10^, Giovanni Galletti^11^, Ian Magill^5^, Iliyan D Iliev^12^, Joonsoo Kang^9^, Jordan Voisine^2^, Josh Choi^5^, Julia Merkenschlager^13^, Jun R. Huh^5^, Katharine Block^7^, Ken Cadwell^10^, Kennidy K. Takehara^11^, Kevin Osum^7^, Laurent Brossay^14^, Laurent Gapin^15^, Liang Yang^5^, Lizzie Garcia-Rivera^1^, Marc K. Jenkins^7^, Maria Brbic^16^, Maria-Luisa Alegre^2^, Marion Pepper^8^, Mariya London^17^, Matthew Stephens^2^, Maurizio Fiusco^16^, Melanie Vacchio^3^, Michael Starnbach^5^, Michel Nussenzweig^13^, Mitch Kronenberg^18^, Myriam Croze^19^, Nalat Siwapornchai^5^, Nathan Morris^12^, Nicole E. Scharping^11^, Nika Abdollahi^19^, Nitya Mehrotra^2^, Odhran Casey^5^, Olga Barreiro del Rio^5^, Paul Thomas^20^, Peter Carbonetto^2^, Remy Bosselut^3^, Rocky Lai^9^, Sam Behar^9^, Sam Borys^14^, Sara E. Hamilton^7^, Sara Mostafavi^8^, Sara Quon^11^, Serge Candéias^21^, Shanelle Reilly^14^, Shanshan Zhang^5^, Siba Smarak Panigrahi^16^, Sofia Kossida^19^, Stefan Muljo^3^, Stefan Schattgen^20^, Stefani Spranger^22^, Steve Jameson^7^, Susan M. Kaech^1^, Takato Kusakabe^12^, Taylor Heim^22^, Tianze Wang^8^, Tomoyo Shinkawa^9^, Ulrich von Andrian^5^, Val Piekarsa^5^, Véronique Giudicelli^19^, Vijay Kuchroo^5^, Woan-Yu Lin^12^, Ziang Zhang^2^

1. NOMIS Center, Salk Institute for Biological Sciences, 2. The University of Chicago, 3. National Institutes of Health, 4. Allen Institute for Immunology, 5. Harvard Medical School, 6. Dept of Dermatology and Immunology, University of Pittsburgh, 7. University of Minnesota, 8. University of Washington, 9. UMass Chan Medical School, 10. University of Pennsylvania, 11. University of California San Diego, 12. Weill Cornell Medicine, 13. The Rockefeller University, 14. Brown University, 15. University of Colorado Anschutz Medical Campus, 16. Swiss Federal Institute of Technology, Lausanne, 17. New York University, 18. La Jolla Institute, 19. IMGT, Univ Montpellier, 20. St. Jude Children’s Research Hospital, 21. Alternative Energies and Atomic Energy Commission, Grenoble, 22. Massachusetts Institute of Technology

---

**Extended Data Figure 1.**
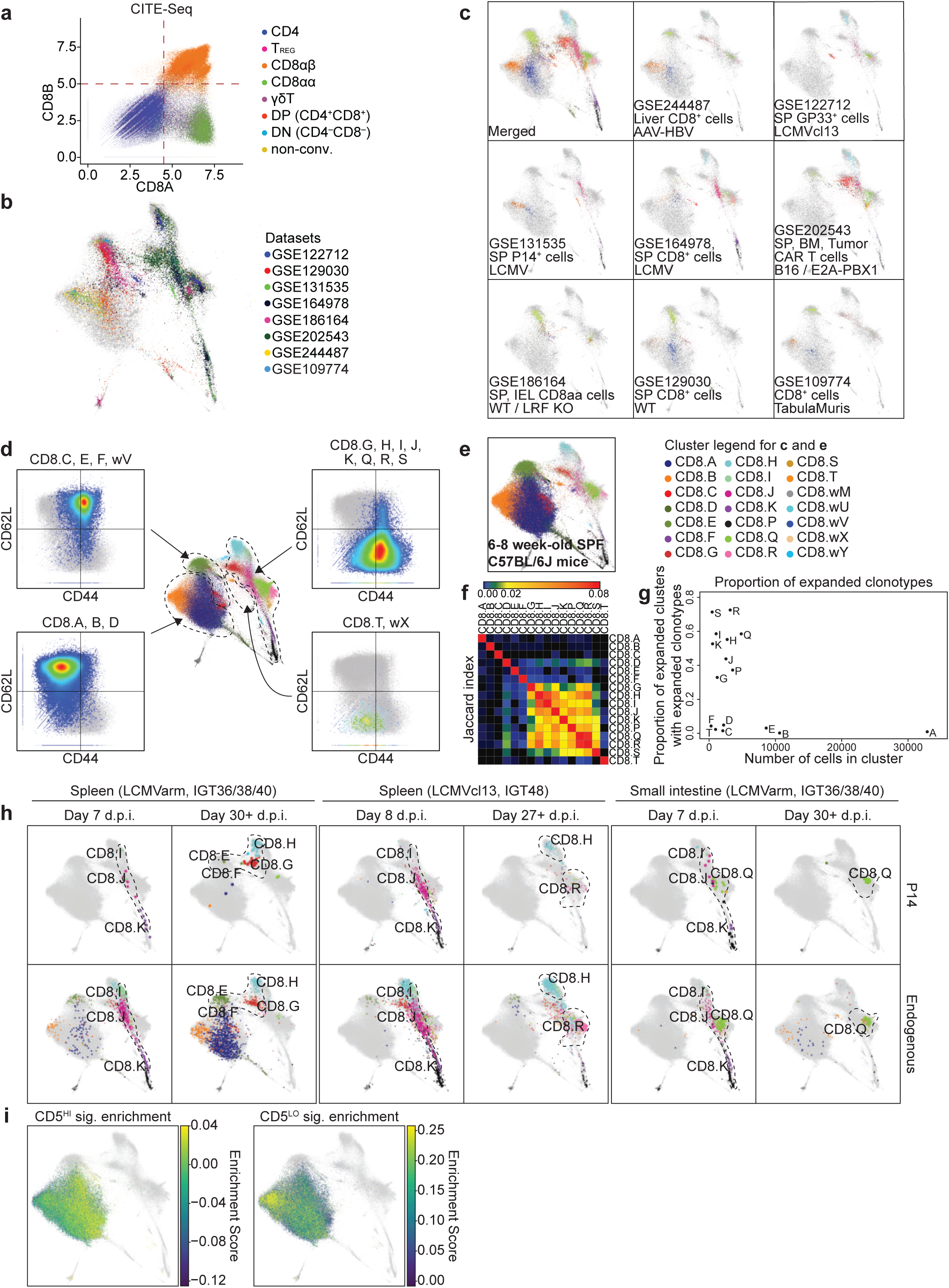
Comprehensive integration of the immgenT-CD8 framework. **a**, Scatter plot showing CITE-seq expression levels of CD8A and CD8B proteins across the full immgenT dataset, showing uniform co-expression of CD8α and CD8β on cells annotated as the CD8αβ^+^ T cell lineage; **b,c**, T-RBI analysis of eight published datasets (see **Methods** and immgenT-Cosmology manuscript^34^) shown together (**b**) or individually (**c**). Tissue of origin, cell type, and condition of each GEO accession code are indicated on each plot (see **Extended Data Table 3**); **d**, CD8-MDE annotated by the 21 clusters, with adjacent scatter plots partitioning the space into four broad areas based on CITE-seq protein levels of CD62L and CD44 (naive/resting: CD62L^+^CD44^−^; effector: CD62L^−^CD44^+^; central-memory-like: CD62L^+^CD44^+^; double-negative: CD62L^−^CD44^−^); **e**, CD8-MDE showing CD8^+^ T cells from healthy 6-8-week-old specific-pathogen-free (SPF) C57BL/6J mice. Cells are colored by cluster identity; background immgenT-CD8 are shown in gray; **f**, Heatmap showing the incidence of clonotype sharing between clusters (Jaccard index); **g**, Scatter plot of the proportion of cells with expanded clonotypes versus cluster size; **h**, CD8-MDE showing LCMV-specific CD8^+^ T cells (P14 transgenic cells, top) and endogenous CD8^+^ T cells (bottom) in the spleen and small intestine of mice infected with LCMV-Armstrong or –Clone 13, at 7 days post-infection (acute phase) or at the late phase (days 27-30+). Cells are colored by cluster identity; background immgenT-CD8 are shown in gray; **i**, CD8-MDE plot showing enrichment of published CD5^HI^ and CD5^LO^ naive T cell signatures^35^ preferentially in clusters A and B, respectively. **Abbreviations**: AAV-HBV, adeno associated virus hepatitis B virus; SP, spleen; BM, bone marrow; CAR, chimeric antigen receptor; B16, B16 melanoma model; E2A-PBX1, E2A-PBX1 leukemic cell line; IEL, intraepithelial lymphocytes; WT, wild-type; LRF KO, Leukemia/lymphoma-related factor knock-out; SPF, specific pathogen-free; d.p.i., days post-infection; sig., signature.

**Extended Data Figure 2.**
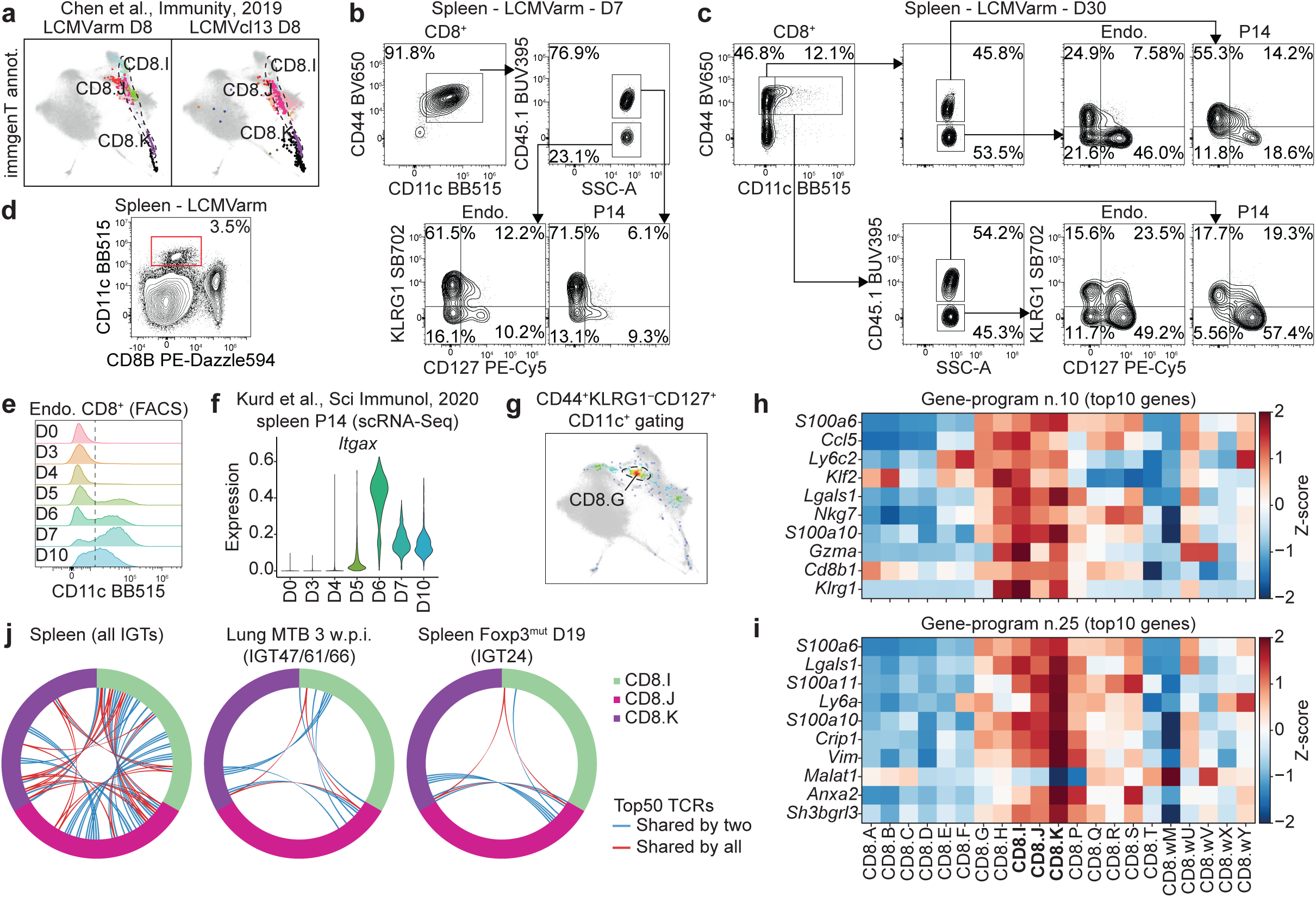
Acute effector CD8^+^ T cell states (clusters CD8.I, .J, and .K). **a**, T-RBI of published datasets of LCMV-specific T cells (P14 transgenic cells) from secondary lymphoid organs at day 8 (acute phase) post LCMV-Armstrong (left) and Clone 13 (right) from Chen et al.^91^. Cells are colored by immgenT cluster identity; background immgenT-CD8 are shown in gray; **b,c**, Flow cytometry plots showing KLRG1 and CD127 expression on splenic CD44^+^CD11c^+^ or CD44^+^CD11c^−^ endogenous and P14 transgenic CD8^+^ T cells at day 7 (**b**) or day 30 (**c**) after LCMV-Armstrong, demonstrating that the CD11c gate specifically captures multiple classical effector subsets; **d**, CD11c expression on CD8B^−^ leukocytes (dendritic cells as positive control) from spleen (LCMV day 7), illustrating lower expression on T cells; **e**, Flow cytometry histograms of CD11c protein over time on splenic endogenous CD8^+^ T cells from day 0 to day 10 post LCMV-Armstrong infection; **f**, *Itgax* (encoding CD11c) transcript expression in splenic P14 CD8^+^ T cells across the same time course from a published scRNA-seq dataset^92^; **g**, CD8-MDE highlighting cells bioinformatically gated as CD44^+^KLRG1^−^CD127^+^CD11c^+^ (common memory-precursor phenotype). These cells show preferential accumulation in cluster G, with only minor representation in other memory clusters; **h,i**, Heatmaps showing expression (Z-score normalized) of the top 10 genes in gene-program GP10 (**h**) and GP25 (**i**) across all immgenT-CD8 clusters; **j**, Circos plots showing the top 50 TCR clones and their sharing among clusters CD8.I, .J, and .K in the spleen (left), in the lung of infected mice with MTB 3 weeks post-infection (middle), and in the spleen of 19-day-old Foxp3-deficient mice at peak of inflammation (right). **Abbreviations**: annot., annotation; D, day; Endo., endogenous; n., number; MTB: Mycobacterium tuberculosis.

**Extended Data Figure 3.**
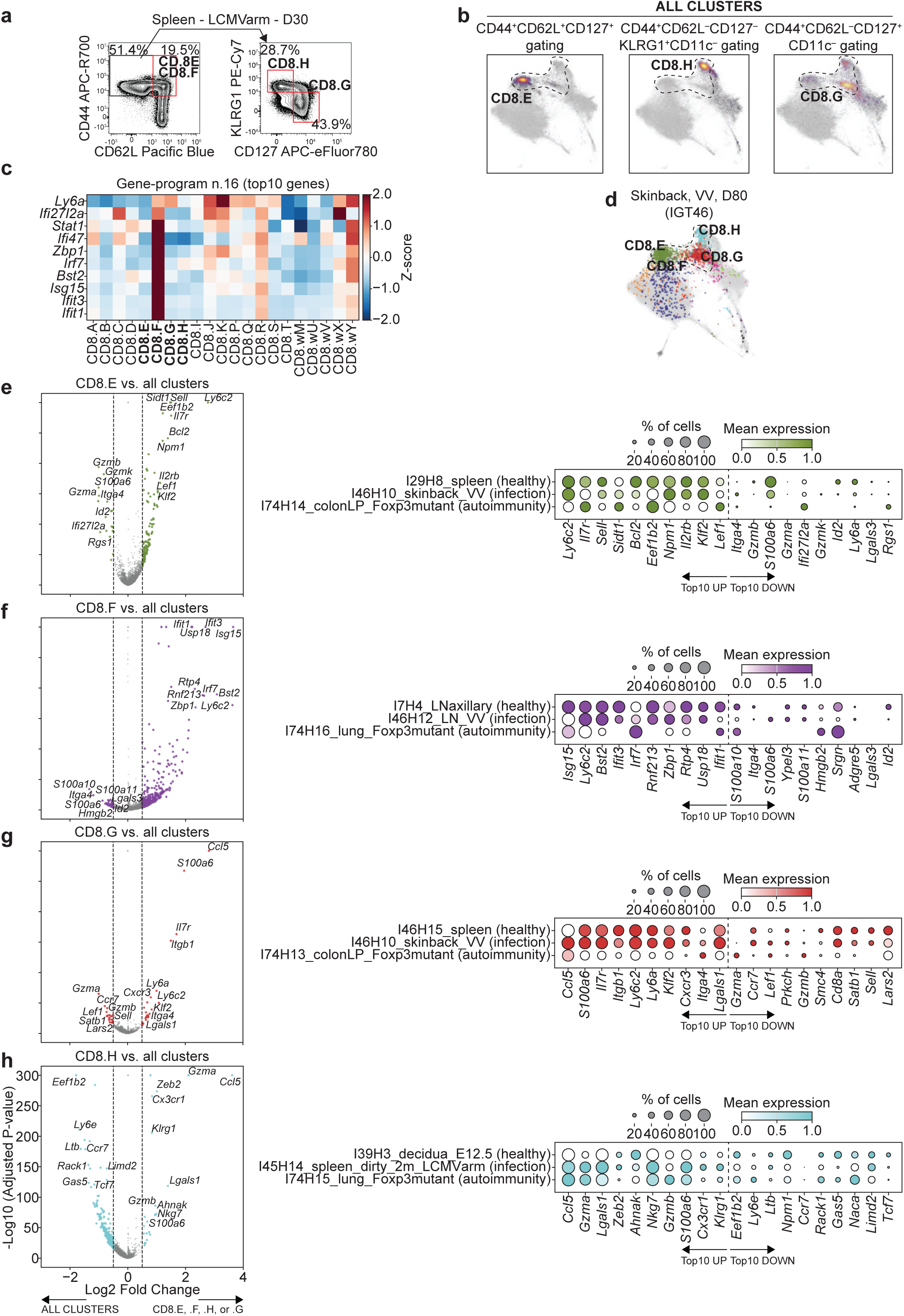
Phenotypic and transcriptional signatures of circulating memory states (clusters CD8.E, .F, .G, and .H). **a**, Representative flow cytometry plots showing the CITE-seq-derived gating strategies shown in Fig. 4c on endogenous CD8^+^ T cells from spleen at day 30 after LCMV-Armstrong infection (percentages shown in gates); **b**, CD8-MDE highlighting cells bioinformatically gated using the CD8.D (left), CD8.H (middle) and CD8.G (right) gating strategies (shown above each plot); **c**, Heatmap showing expression (Z-score normalized) of the top 10 genes from interferon-responsive GP16 across clusters. Cluster CD8.F is selectively enriched for this program; **d**, CD8-MDE showing CD8^+^ T cells from the skin at day 80 after vaccinia virus scarification upon 1-fluoro-2,4-dinitrobenzene (DNFB) challenge^51^, showing bystander-activated CD8^+^ T cells mapping to clusters CD8.E-H; **e-h**, Volcano plots showing gene signatures for CD8.E (**e**), .F (**f**), .G (**g**), and .H (**h**) compared with all other clusters (left panels). Dot plots showing the mean expression (column-scaled) of the top 10 up– or down-regulated genes in each signature across representative samples (right panels). **Abbreviations**: D, day; n., number; VV, vaccinia virus; vs., versus.

**Extended Data Figure 4.**
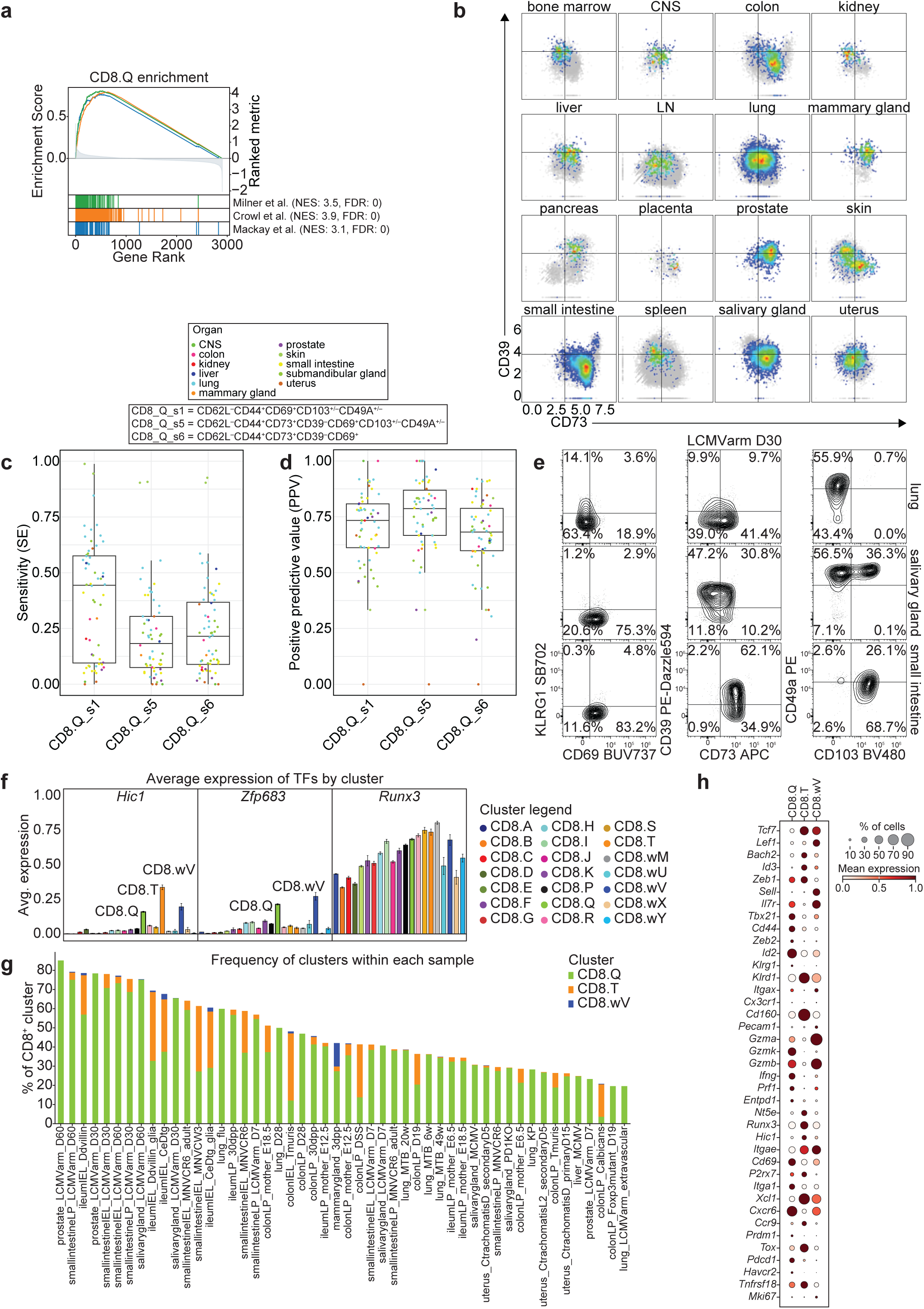
Surface-marker and transcriptional heterogeneity of tissue-resident memory CD8^+^ T cells in cluster CD8.Q. **a**, Gene Set Enrichment Analysis (GSEA) of published T_RM_ signatures^6,52,53^ using the CD8.Q gene signature; **b**, Scatter plots showing CITE-seq protein expression of CD39 and CD73 within CD8.Q cells across tissues. Cells are colored by density; background immgenT-CD8 are shown in gray; **c,d**, Box plots of sensitivity (**c**) and positive predictive value (**d**) for different CITE-seq-guided gating strategies for CD8.Q across samples (see also **Extended Data Table 6**); **e**, Flow cytometry plots showing the expression of CD69, KLRG1, CD39, CD73, CD103, and CD49A in CD44^+^ intravenous-label-negative P14 CD8^+^ T cells from lung, salivary gland, and small intestine at memory time points after LCMV-Armstrong; **f**, Bar plot showing average expression of selected T_RM_-associated transcription factors (*Hic1*, *Zfp683*, *Runx3*) across all immgenT-CD8 clusters; **g**, Stacked bar plot showing the proportion of clusters CD8.Q, CD8.T, and CD8.wV within the top 50 samples. Only samples with at least 10 total CD8^+^ T cells recovered in at least one cluster are shown; **h**, Dot plot showing scaled mean expression of selected genes across clusters CD8.Q, CD8.T, and CD8.wV. **Abbreviations**: sig., signature; CNS, central nervous system; LN, lymph node; s, strategy; D, day; TF, transcription factor.

**Extended Data Figure 5.**
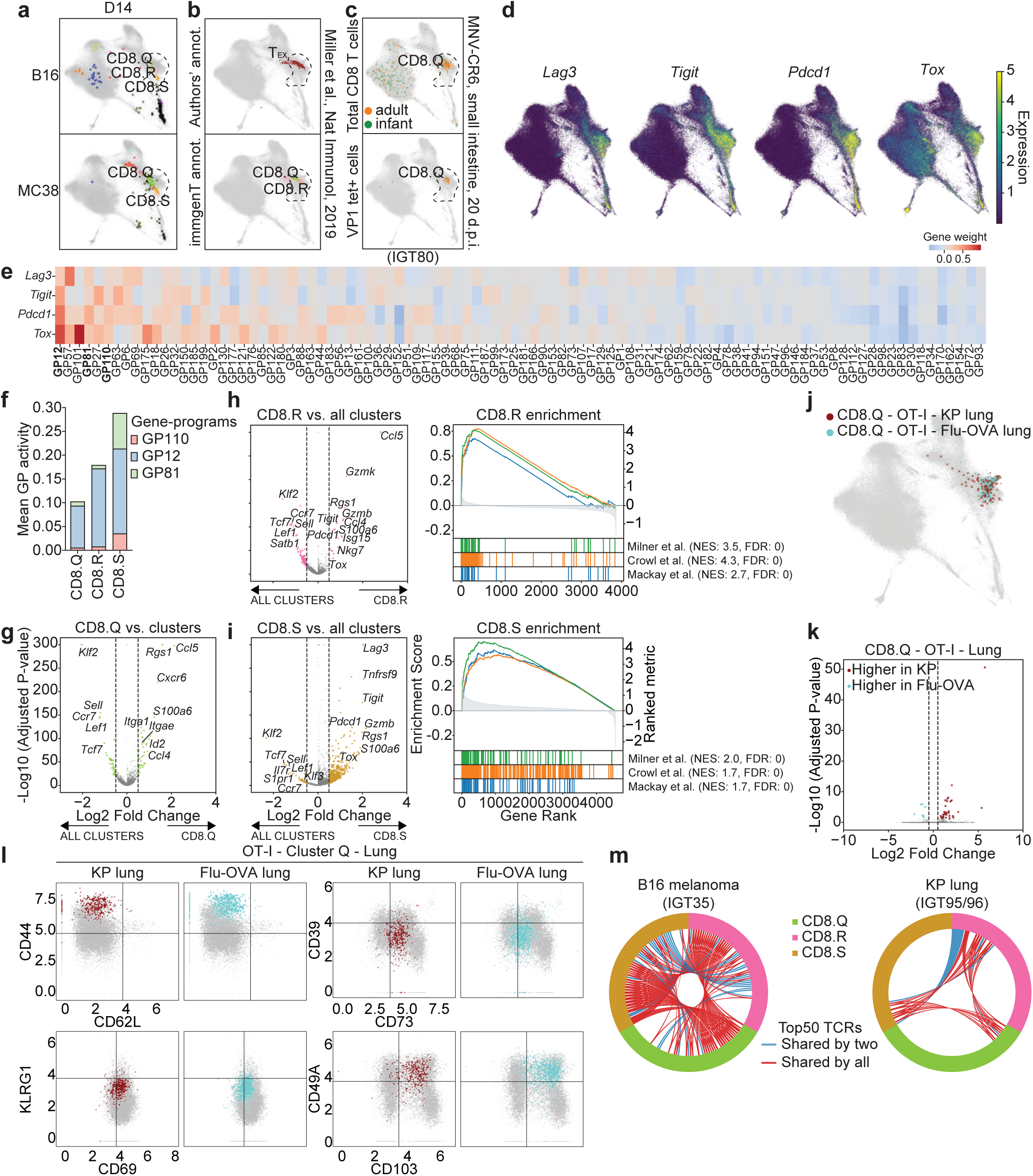
Detailed mapping of the exhaustion/residency continuum and progenitor-exhausted cells. **a**, T-RBI of published datasets of cells from B16 melanoma and MC38 (GSE316401) at day 14 post-implantation. Cells are colored by immgenT cluster identity; background immgenT-CD8 are shown in gray; **b**, T-RBI of LCMV-specific cells in mice chronically infected with LCMV-Clone 13 on day 28 post infection from Miller et al.^64^. Only exhausted T_EX_ cells, as annotated by the authors, are shown. Cells are colored by the authors’ annotation (top) and immgenT cluster identity (bottom); background immgenT-CD8 are shown in gray; **c**, CD8-MDE showing CD8^+^ T cells isolated from the small intestine of mice chronically infected with murine norovirus strain CR6 (MNV-CR6). Cells are colored by adult (orange) or infant (green); **d**, Feature plots showing expression of common exhaustion-associated genes (*Lag3*, *Tigit*, *Pdcd1*, *Tox*); **e**, Heatmap showing gene score for *Lag3*, *Tigit*, *Pdcd1*, and *Tox* across gene-programs; **f**, Stacked bar plot showing the activity of exhaustion-like gene-programs GP12, GP81, and GP110 in clusters CD8.Q, .R, and .S; **g**, Volcano plots showing the gene signature of CD8.Q (versus all other clusters); **h,i**, Volcano plots showing the gene signature of CD8.R (**h**) and S (**i**) (versus all other clusters) (left) with Gene Set Enrichment Analysis (GSEA) of published T_RM_ signatures^6,52,53^ (right); **j**, CD8-MDE showing flu-specific cells and tumor-specific cells (both OT-I transgenic) from the lung of mice infected with flu-OVA (light blue) or implanted with KP lung tumors (dark red) mapping to CD8.Q; **k**, Volcano plot of differentially expressed genes in CD8.Q OT-I cells from KP lung tumor (dark red) versus flu-OVA-infected lung (light blue); **l**, Scatter plots showing CD62L, CD44, CD69, KLRG1, CD39, CD73, CD103 and CD49A expression in CD8.Q OT-I cells from KP lung tumor (dark red) and flu-OVA-infected lungs (light blue) (CITE-seq; log1p(CP10K)); **m**, Circos plots showing the top 50 TCR clones and their sharing among clusters CD8.Q, CD8.R, and CD8.S in B16 melanoma (left) and KP lung cancer (right). **Abbreviations**: D, day; GP, gene-program; vs., versus; NES, normalized enrichment score; FDR, false discovery rate.

**Extended Data Figure 6.**
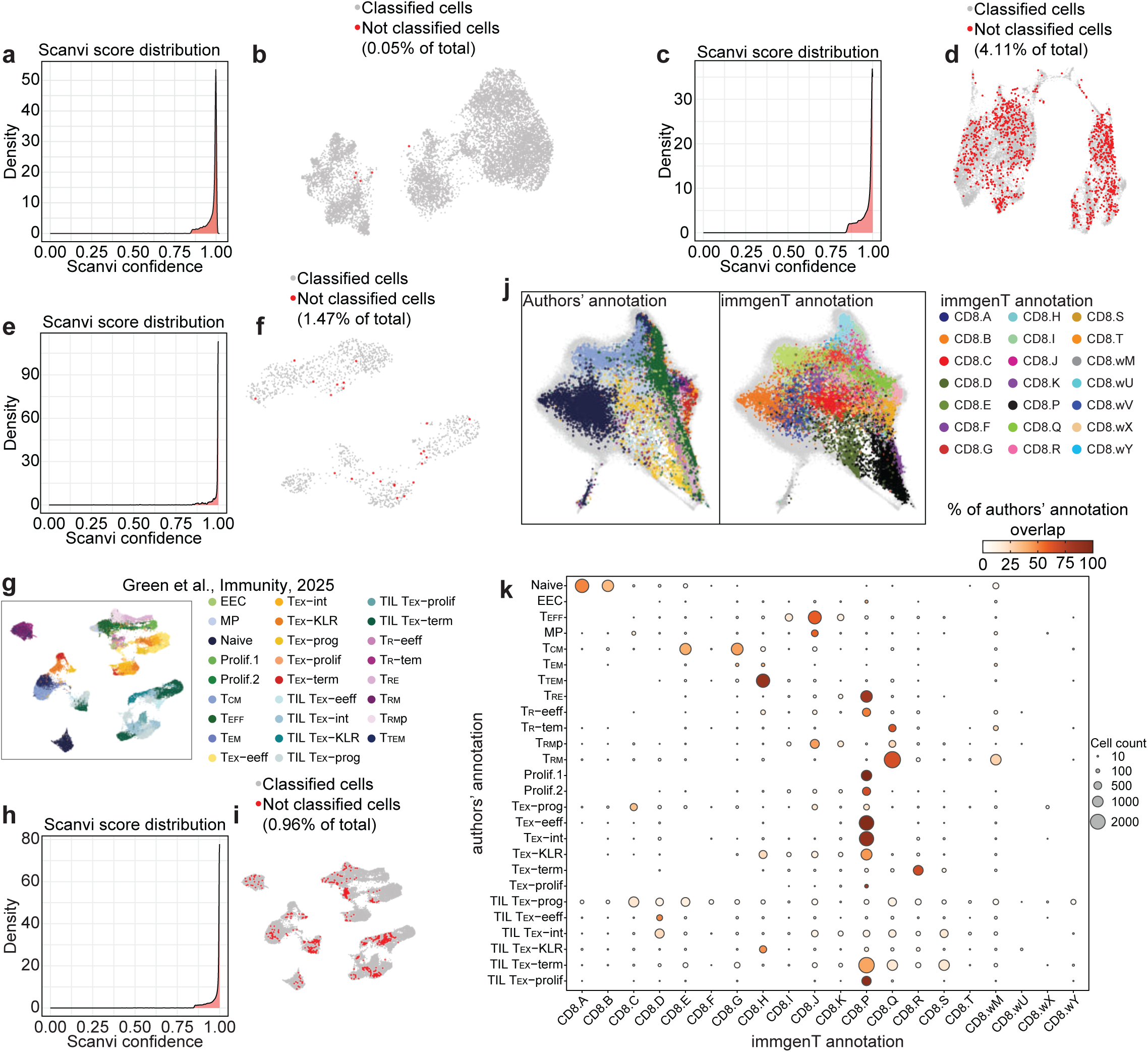
Performance and practical implementation of the T-RBI analysis. **a,b**, T-RBI statistics of the AAV-HBV CD8^+^ T cell dataset by Conceição-Neto et al.^68^. Histogram of scANVI confidence scores for immgenT cluster assignment (**a**) and UMAP showing the immgenT unclassified cells (red) intermingled with annotated cells (gray) (**b**). These cells did not form coherent novel clusters; **c,d**, T-RBI statistics of the CAR T cell dataset by Zhu et al.^69^. Histogram of scANVI confidence scores for immgenT cluster assignment (**c**) and UMAP showing the immgenT unclassified cells (red) intermingled with annotated cells (gray) (**d**). These cells did not form coherent novel clusters; **e,f**, T-RBI statistics of the primary biliary cholangitis versus wild-type dataset by Han et al.^70^. Histogram of scANVI confidence scores for immgenT cluster assignment (**e**) and UMAP showing the immgenT unclassified cells (red) intermingled with annotated cells (gray) (**f**). These cells did not form coherent novel clusters; **g**, UMAP projection of the scRNA-Seq layer from Green et al. (2025)^71^ showing P14 CD8^+^ T cells across LCMV-Armstrong, LCMV-Clone 13, and four GP33-expressing tumor models. Cells colored by the authors’ annotations; **h,i**, T-RBI statistics of the P14 dataset by Green et al.^71^. Histogram of scANVI confidence scores for immgenT cluster assignment (**h**) and UMAP showing the immgenT unclassified cells (red) intermingled with annotated cells (gray) (**i**). These cells did not form coherent novel clusters; **j**, The same cells projected onto the immgenT CD8-MDE (gray background) using T-RBI and colored by authors’ original annotations (left) or immgenT cluster assignment (right); **k**, Dot plot showing the distribution of the authors’ original clusters across immgenT clusters (see **Extended Data Table 9**).

**Extended Data Figure 7.**
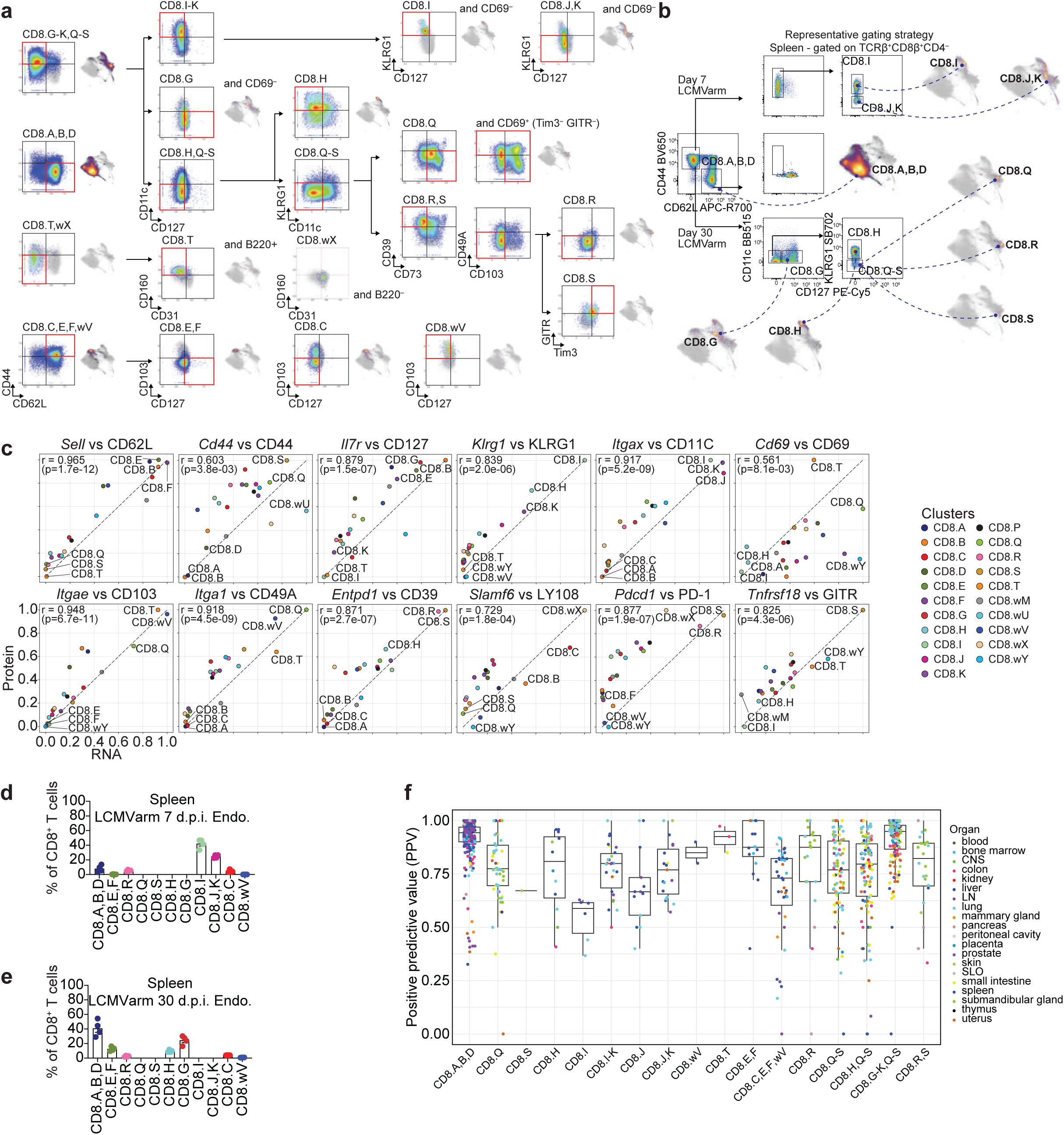
immgenT-CD8 flow cytometry panel and validation. **a**, Scatter plots showing CITE-seq-based gating strategies for all 21 clusters using the immgenT-CD8 panel. CD8-MDE plots highlight the cells in the successive gated red boxes; **b**, Representative flow cytometry gating strategy and CD8-MDE plots illustrating how to isolate the main immgenT-CD8 clusters; **c**, Scatter plots showing mean RNA (log1p normalized counts) versus surface protein (log1p normalized counts) expression for the indicated gene-protein pairs across the 21 CD8^+^ clusters. Spearman correlation coefficients were calculated on the raw cluster means. For visualization, RNA and protein values were min-max scaled independently within each panel. The dashed diagonal line indicates perfect concordance; **d,e**, Bar plot showing frequencies of clusters recovered by flow cytometry using the immgenT-CD8 panel on day 7 (**d**; see Fig. 3b**-e**) and day 30 (**e**; see Fig. 4c and **Extended Data** Fig. 3a**,b**) post LCMV-Armstrong; **f**, Box plots of positive predictive value for the immgenT-CD8 panel gating strategies across clusters (**Extended Data Table 4**). **Abbreviations**: d.p.i., days post-infection.

## EXTENDED DATA TABLE LEGENDS

**Extended Data Table 1. immgenT-CD8 samples.**

**Extended Data Table 2. Summary of endogenous and antigen-specific cell counts across samples, conditions and tissues (only samples with at least 10 total CD8^+^ T cells recovered in at least one cluster are shown).**

**Extended Data Table 3. List of external published CD8^+^ T cell datasets projected onto the immgenT reference using T-RBI.**

**Extended Data Table 4. Performance metrics (sensitivity, specificity, PPV, and NPV) for CITE-seq-derived gating strategies across CD8^+^ T cell states.**

**Extended Data Table 5. Correspondence between immgenT clusters and common annotations.**

**Extended Data Table 6. CITE-seq-guided flow-cytometry-like gating strategies for identifying CD8.Q T cell state.**

**Extended Data Table 7. Distribution of original cluster annotations from a published dataset of intrahepatic CD8^+^ T cells in an HBV mouse model across immgenT-CD8 T cell clusters.**

**Extended Data Table 8. Distribution of original cluster annotations from a published CAR T cell dataset across immgenT-CD8 T cell clusters.**

**Extended Data Table 9. Distribution of original cluster annotations from a published infection/tumor P14 CD8^+^ T cell dataset across immgenT-CD8 T cell clusters.**

**Extended Data Table 10. Flow cytometry panel for discrimination of the main immgenT-CD8 T cell states. Extended Data Table 11. CD8^+^ cluster signatures.**

**Extended Data Table 12. List of gene-programs analyzed in this study.**

## Notes

https://www.immgen.org/ImmGenT/

https://www.ncbi.nlm.nih.gov/geo/query/acc.cgi?acc=GSE297097

https://www.ncbi.nlm.nih.gov/geo/query/acc.cgi?acc=GSE316401

https://doi.org/10.5281/zenodo.21839962

### Competing Interest Statement

The authors have declared no competing interest.

### Summary of Updates

Edits to text, main and extended data figures.

